# Identification of potential sepsis therapeutic drugs using a zebrafish rapid screening approach

**DOI:** 10.1101/2024.10.09.617431

**Authors:** Mark Widder, Chance Carbaugh, William van der Schalie, Ronald Miller, Linda Brennan, Ashley Moore, Robert Campbell, Kevin Akers, Roseanne Ressner, Monica Martin, Michael Madejczyk, Blair Dancy, Patricia Lee, Charlotte Lanteri

## Abstract

In the military, combat wound infections can progress rapidly to life-threatening sepsis. Discovery of effective small molecule drugs to prevent and/or treat sepsis is a priority. To identify potential sepsis drug candidates, we used an optimized larval zebrafish model of endotoxicity/sepsis (Philip et al., 2017) to screen commercial libraries of U.S. Food and Drug Administration (FDA)- approved drugs and other active pharmaceutical ingredients (API) known to affect pathways implicated in the initiation and progression of sepsis in humans (i.e., inflammation, mitochondrial dysfunction, coagulation, and apoptosis). We induced endotoxicity in 3- and 5-day post fertilization larval zebrafish (characterized by mortality and tail fin edema (vascular leakage)) by immersion exposure to *Pseudomonas aeruginosa* 60 μg/mL lipopolysaccharide (LPS) for 24 hours, then screened for the rescue potential of 644 selected drugs simultaneously with LPS at 10 μM. After LPS exposure, we used a neurobehavioral assay (light-dark test) to further evaluate rescue from endotoxicity and to determine possible off-target drug side effects. We identified 29 drugs with > 60% rescue of tail edema and mortality. Three drugs (Ketanserin, Tegaserod, and Brexpiprazole) produced 100% rescue and did not differ from the controls in the light-dark test, suggesting a lack of off-target neurobehavioral effects. Further testing of these three drugs at a nearly 100% lethal concentration of *Klebsiella pneumoniae* LPS (45 μg/mL) showed 100% rescue from mortality and 88%-100% mitigation against tail edema. The success of the three identified drugs in a zebrafish endotoxicity/sepsis model warrants further evaluation in mammalian sepsis models.

## 1. Introduction

The emergence of multidrug-resistant (MDR) bacteria is a global problem for current antibiotic therapies, a difficult challenge for hospitals worldwide, and a threat to effective treatment of infections among combat-wounded Service Members. Combat wound infections can progress rapidly to life-threatening sepsis with a median time of three days from injury to the onset of sepsis (Weintrob et al., 2018). A total of 5,278 sepsis hospitalizations were recorded in the active component of the U.S. military from 2011 to 2020 (Snitchler et al., 2021). A recent review of the Department of Defense Trauma Registry from 2007 to 2020 found that patients with combat injury related wound infections who were diagnosed with sepsis had significantly higher mortality rates than non-septic patients (21.9% vs. 4.3%, p < 0.001) (Carius et al., 2021).

The discovery and development of therapeutics that mitigate against host-directed mechanisms responsible for triggering sepsis, to be administered in combination with antibiotics, is a viable research and development path for improving options available for preventing morbidity and mortality associated with sepsis. Sepsis is caused by a dysregulated host response to infection involving a complex (and not entirely understood) pathophysiology that leads to life-threatening organ dysfunction (Schlapbach et al. 2020). Because the mortality rate of sepsis patients continues to be comparatively high relative to non-septic patients, there is an urgent need for improved sepsis therapeutics that specifically address systemic infection and sepsis remediation, which can include modulation of the host response. Small molecule drugs that mitigate against dysregulated host responses to bacterial infection may provide the required systemic approach needed to prevent or treat sepsis.

One problem with identifying new drugs or drug combinations for wound infection and sepsis is the lack of a preclinical model that has better predictive capability than cell-based systems but with higher throughput and lower cost than mammalian models (Philip et al., 2017). A whole animal model such as the zebrafish allows determination of drug bioactivity, toxicity, and off-target side effects early in the drug discovery process (Bowman and Zon, 2010), thus filling a gap between in vitro or in silico testing and pre-clinical rodent testing. Zebrafish models of sepsis offer an opportunity for establishing a low-cost and rapid screen to accelerate discovery of potential new small molecule therapies to prevent or treat sepsis. Pathophysiological symptoms observed in septic humans, such as dysregulated inflammatory responses (cytokine storms), tachycardia, endothelial leakage, and progressive edema have been observed in zebrafish infected with bacteria (Barber et al., 2016). Zebrafish have a high degree of genetic homology with humans (Howe et al., 2013) and have been used as a model for a range of human diseases (Lieschke and Currie, 2007). Zebrafish have demonstrated success in rapid drug screening studies (Fleming and Alderton, 2013); Patton et al. (2021) note that “Numerous drug treatments that have recently entered the clinic or clinical trials have their genesis in zebrafish.”

Zebrafish are well suited for studies of bacterial infection and sepsis. Zebrafish have been widely used for studying host-pathogen interactions and have an innate and adaptive immune system that become active at about two days and three weeks post-fertilization, respectively (Philip et al., 2017). The zebrafish immune system has many similarities to mammals (van der Sar, 2004; Sullivan and Kim, 2008; Allen and Neely, 2010; Masud et al., 2017; Torraca and Mostowy, 2018; Sisodia et al., 2020; Gomes and Mostowy, 2020), and zebrafish have been used in infection models for bacterial species of particular concern to the military, e.g., *Klebsiella pneumoniae* (Marcoleta et al., 2018; Leber, 2019; Zhang et al., 2019) and *Pseudomonas aeruginosa* (Clatworthy et al., 2009; Sullivan et al., 2017; Nogaret et al., 2021). In both mammals and zebrafish, the lipopolysaccharide (LPS) component of the cell wall in Gram-negative bacteria can trigger inflammatory responses (Forn-Cuní et al., 2017).

To identify potential sepsis therapeutic drugs, a zebrafish endotoxicity/sepsis model (Philip et al., 2017) was first optimized by evaluating the best combination of endpoints (mortality, tail edema, and reactive oxygen species (ROS) production), LPS sources and concentrations, and exposure times to evaluate the potential of the test drugs to provide rescue from sepsis symptoms. After testing with model compounds previously shown to have efficacy in the zebrafish endotoxicity model, the optimized larval zebrafish model was used to screen the rescue potential of 644 compounds known to affect pathways implicated in the initiation and progression of sepsis in humans. These compounds were derived from commercially available libraries identified to have already been approved by the U.S. Food and Drug Administration (FDA) or other international regulatory agencies for other purposes. Following LPS exposure, a neurobehavioral assay (light-dark test) was used to determine possible off-target drug side effects. Top rescue drugs were tested at a nearly 100% lethal concentration of *Klebsiella pneumoniae* LPS (45 μg/mL) to further evaluate rescue potential. The drug candidates with the best ability to rescue zebrafish larvae from LPS-induced endotoxicity without neurobehavioral effects are candidates for further evaluation in mammalian sepsis models.

## 2. Materials and methods

### 2.1. Zebrafish husbandry

Adult *Danio rerio* (zebrafish AB strain) from in-house breeding stocks were used to produce high quality embryos used for testing. The zebrafish stocks are outcrossed periodically with new stocks of AB zebrafish obtained from the Zebrafish International Resource Center (Eugene, OR, USA). Zebrafish were maintained at an Association for Assessment and Accreditation of Laboratory Animal Care International (AAALAC) approved facility at the Walter Reed Army Institute for Research (WRAIR). All experiments were performed in compliance with AAALAC guidelines. The zebrafish colony was maintained on a 14-h light and 10-h dark photoperiod under full spectrum LED lighting. The adult zebrafish colony was housed in either flow-through aquaculture racks or in tanks with these water quality conditions: water temperature 27.5 ± 1.5 °C; dissolved oxygen 7.52 ± 0.3 mg/L; pH 7.56 ± 0.3; alkalinity 110 to 180 mg/L as CaCO_3_; hardness 150 to 210 mg/L as CaCO_3_; conductivity 651 ± 75 μS/cm; and total ammonia less than 0.1 mg/L as NH_3_. Adult zebrafish were fed three times daily on weekdays: 2 feedings of Gemma Micro 300 (Skretting Zebrafish, Westbrook, ME) and 1 feeding of live brine shrimp nauplii (Brine Shrimp Direct, Ogden, UT). On weekends, adult zebrafish received 2 feedings: one Gemma Micro 300 and one live brine shrimp nauplii. Zebrafish aged 6–18 months were bred in I-SPWAN-S breeding chambers (Techniplast, West Chester, PA) to supply embryos for drug screening. Zebrafish embryos from the spawning events were collected in glass petri dishes containing freshly prepared embryo media (EM, see supplemental data for formulation). At the conclusion of the study, all surviving larvae were euthanized with sodium hypochlorite according to the *AVMA Guidelines for Euthanasia of Animals, 2020*.

### 2.2. Zebrafish model optimization: LPS and endpoint screening

The endotoxicity/sepsis zebrafish model used was based on the method of Philip et al. (2017). In this model, larval zebrafish (3 days post fertilization (dpf)) were exposed to *Escherichia coli* 0111:B4 LPS by immersion in microplate wells for 24 hours, and three endpoints were measured: mortality, tail edema, and ROS production. To facilitate rapid and repeatable drug screening for rescue from LPS-induced endotoxicity, the responsiveness of these three endpoints to alternative LPS sources were determined.

Three sources of LPS were evaluated: *E. coli* 0111:B4, *P. aeruginosa* Sigma L9143, and *K. pneumoniae. E. coli* 0111:B4 LPS was selected as the reference LPS used by Philip et al. (2017). *P. aeruginosa* LPS is commonly associated with infections in hospitals, accounting for an estimated 32,600 cases and resulting in an estimated 2,700 deaths in 2017 (CDC Report 2019). *P. aeruginosa* infections are also a common cause of sepsis in both patients with severe burns and immunocompromised patients. *K. pneumoniae* LPS has military relevance related to battlefield injury infections (Kiley et al., 2021). Initial range finding LC50 tests with larval zebrafish were conducted for each LPS strain at both 3 dpf and 5 dpf. For the 5 dpf LC50 range find test, 15 zebrafish embryos were exposed in 6 well microplates containing 3.5 mL of 0, 25, 50, 100, 150, and 200 μg/mL of LPS in EM. (*E. coli* 0111:B4 was also tested at higher concentrations, 400, 800, and 1000 μg/mL due a lack of toxicity within the initial dose range.) Well plates were then covered with parafilm and placed into an incubator at 28.5°C for 24 hours. After 6 and 24 hours, the plates were removed and scored for mortality, tail edema, and ROS production. The tail edema endpoint evaluates vascular leakage from caudal fin in response to LPS exposure (Figure 2). ROS production was determined as described by Philip et al. (2017) by adding 2,7-dichloroflurescein diacetate (100μM) to the wells containing the zebrafish larvae and incubating them in a 100% dark photoperiod incubator at 28.5°C for one hour. Florescent images of the incubated zebrafish were then captured with a Keyence BZ-X710 fluorescent microscope using 10X objective with a GFP filter and fixed 1/15 second exposure for all imaging. The 3 dpf LC50 range find was the same as the 5dpf range find with the following changes: 96 well plates were used in place of the 6 well plates, the number of zebrafish was increased from 15 to 32, and the dose ranges tested for *P. aeruginosa* and *K. Pneumoniae* were changed to 10, 20, 30, 40, 50,60, 70, 80, 90, and 100 μg/mL due to the steep mortality curve in the 5 dpf range find. Results from the LPS toxicity testing and endpoint screening were used to develop the optimized testing procedure through evaluation in selected model compounds, as described in the next section and in the results.

### 2.3. Model compound testing

The optimized zebrafish sepsis model was evaluated against compounds previously shown to have efficacy in previous zebrafish sepsis models: fasudil, hydrocortisone, dexamethasone, and fisetin. Fasudil, an anti-vascular leakage compound (inhibitor of the RhoA/Rho-kinase pathway) was shown by Philip et al. (2017) to rescue zebrafish from LPS-induced mortality, ROS production, and tail fin edema. Fasudil also rescued LPS-induced vascular leakage in murine models of experimental sepsis (Li et al., 2010) and reduced acute lung injury in septic rats through inhibition of the Rho/ROCK signaling pathway (Wang et al., 2018). Hsu et al. (2018) found that the corticosteroids hydrocortisone and dexamethasone produced responses similar to a protein tyrosine phosphatase inhibitor (Shp2 (Ptpn11a) inhibitor 11a-1), leading to decreased inflammatory cytokines, increased *il-10* anti-inflammatory activity, reduced tissue damage and preservation of vascular junction proteins in their zebrafish model. Fisetin is a dietary flavonoid that showed inhibition of LPS-induced inflammation and reduced mortality from endotoxic shock in zebrafish larvae; possible mechanisms include crosstalk between GSK-3β/β-catenin and the NF- κB signaling pathways (Molagoda et al., 2021). Endpoints evaluated with model compounds included mortality, tail edema, and ROS at 6- and 24-hours post exposure to varying dose ranges of LPS.

### 2.4. Custom drug library selection

Four commercially available drug discovery libraries including nearly 2500 compounds were selected to evaluate drug targets known to affect pathways implicated in the initiation and progression of sepsis in humans (Adadey et al. (2018), Bergmann et al. (2021), Dare et al. (2009), Huerta and Rice (2019)), including inflammation, mitochondrial dysfunction, coagulation, and apoptosis. Libraries selected included the Selleckchem Cytokine Inhibitor Library (L9500), the Selleckchem Highly Selective Inhibitor Library (L3500), the MedChemExpress Mitochondria-Targeted Compound Library (HY-L089), and the MedChemExpress Coagulation and Anti-coagulation Compound Library (HY-L136). To reduce the high costs associated with developing new or novel drugs, this initial set of candidate compounds was cross-referenced with Selleckchem FDA-Approved Drug Library (L1300) and MedChemExpress FDA-Approved Drug Library (HY-L022). This evaluation identified 644 drugs for sepsis efficacy screening in the optimized zebrafish endotoxicity/sepsis model.

### 2.5. Initial drug screening

Healthy, normally developing embryos collected from the adult AB zebrafish spawning events were treated with 75 μM of 1-phenyl 2-thiourea (PTU) in EM prior to 24 hours post fertilization (hpf) to block pigmentation formation (Karlsson *et al*, 2001). These embryos were then placed into an incubator at 28.5°C until they reached 3 dpf. Embryos were then removed from the incubator and plated individually into the wells of 96 well plates containing 50 μL of EM. A super stock of 120 μg/mL of *P. aeruginosa* LPS was made up in EM in a glass container. This stock was then used to make sub stocks of 120 μg/mL *P. aeruginosa* LPS (positive control) and 120 μg/mL *P. aeruginosa* LPS containing 20 μM of test drug in glass scintillation vials. Immediately after the sub stocks were made, the wells were dosed with either 50 μL of 120 μg/mL of *P. aeruginosa*, 120 μg/mL of *P. aeruginosa* with 20 μM of test drug, or EM (1:1 Dilution). Each 96 well plate had one LPS positive control (n=16), 4 different test drugs with *P. aeruginosa* LPS (n=16 per drug), and one negative control (n=16). Once the plates were dosed, they were covered with parafilm and placed into an incubator at 28.5°C. The plates were removed at 6- and 24-hours post exposure to be scored for tail edema and mortality. All 644 drugs evaluated for rescue from mortality and tail edema induced by *P. aeruginosa* LPS exposure as compared to LPS-treated controls after 24 hours of exposure.

### 2.6. Top rescue drug confirmatory testing and down selection

To further down select the top rescue drugs, only drugs that had >60% rescue of mortality and tail edema induced by *P. aeruginosa* LPS exposure from the initial screen were further evaluated in confirmatory testing. Confirmatory testing followed the same methods as the initial testing but used 5 dpf larvae instead of the 3 dpf larvae that were used for the initial screening because they were easier to handle and replicated the methods of Philip et al. (2017). The 5 dpf larvae have more fully developed organ functionality than 3 dpf larvae, and in a meta-analysis of acute toxicity data for 600 chemicals, Ducharme et al. (2015) found that larval zebrafish acute toxicity data were predictive of rodent and rabbit acute toxicity for 4 and 5 dpf larvae, but not for 3 dpf larvae.

In a second stage of confirmatory testing, drugs that had >80% rescue of mortality and tail edema with the 5 dpf larvae were evaluated further. First, a concentration range find test was used to determine the potential therapeutic window for the drugs. Briefly, 5 dpf zebrafish embryos were placed into the wells of the 96 well plate (one embryo per well), with each well containing 50 μL of EM. Then, an additional 50 μL of either 120 μg/mL of *P. aeruginosa* or 120 μg/mL of *P. aeruginosa* with the test drug (1:1 dilution) was added to the 96 well plate. Each 96 well plate had one positive control (n=16) and 4 different concentrations of the test drug with *P. aeruginosa* LPS (n=16 per concentration). Once the plates were dosed, they were covered with parafilm and placed into an incubator at 28.5°C. The plates were removed at 6- and 24-hours post exposure to be scored for tail edema and mortality. The drug concentrations tested were 10, 1, 0.1, 0.01 and 0.001 μM.

Drugs identified with >80% rescue were subjected to additional testing. Drug pretreatment for 24 hours prior to LPS challenge was evaluated to determine if any prophylactic benefit could be realized with any of the top rescue drugs. The intent was to determine if drug pre-treatment might be used at the point of injury as a preventative therapeutic before the onset of infection and development of sepsis. Parallel groups of fish received a 24-hour drug pre-treatment or no drug pre-treatment prior to LPS challenge. For this test, 4 dpf PTU-treated zebrafish embryos were placed into 96 well plates containing 100 μL of EM. The 24-hour drug pre-treatment group had a 10 μM dose of test drug added when the zebrafish embryos were 4 dpf to allow for a 24-hour pre-treatment prior to LPS exposure at 5 dpf. A parallel group that did not receive 24-hour drug pre-treatment was evaluated to provide comparative results due to the slight change in LPS challenge/toxicity required to complete the studies. All plates were covered with parafilm and placed into an incubator at 28.5 °C for 24 hours. After 24 hours, all treatments had an additional 100 μL of either 120 μg/mL of *P. aeruginosa*, 120 μg/mL of *P. aeruginosa* with test drug (to maintain a 10 μM final in well drug concentration and 60 μg/mL of *P. aeruginosa* LPS), or EM alone (control group). Once the plates were dosed, they were covered with parafilm and placed into an incubator at 28.5°C. The plates were removed at 6- and 24-hours post exposure and scored for tail edema and mortality. Once the 24-hour tail edema and mortality endpoints were collected, a neurobehavioral test was conducted as described in section 2.8.

The top 3 drugs (Ketanserin, Tegaserod, and Brexpiprazole) identified from the confirmatory testing were also tested against another strain of LPS to determine if the candidate drug showed efficacy in more than one LPS strain. These tests followed the same methods described in the 5 dpf zebrafish confirmatory testing, but *K. pneumoniae* LPS (45 μg/mL final in well concentration) was used instead of *P. aeruginosa* LPS.

### 2.7. Neurobehavioral testing

All groups were evaluated at the end of the study using a neurobehavioral assay to evaluate deviations from normal fish behavior and further evaluate rescue from sepsis, as well as to determine if any of the drugs might be exhibiting off-target side effects. This assay is a modified version of the larval zebrafish light dark transition assay with an added UV light (wavelength = 295 nm) phase to capture any compound specific neurological effects to UV light. (Basnet et al., 2019, Kokel et al., 2013, Peng et al., 2016). The light period assesses deviations from normal locomotor behavior while the rapid transition to dark provides a measure of stress response by identifying any deviations from the normal heightened locomotor activity caused by the transition to dark. The neurobehavioral assay used a custom trial control script created by Noldus Ethovision XT 15, and it was executed in a Noldus DanioVision system. Zebrafish larvae in microplates were allowed to habituate in the DanioVision system for 5 minutes under white light conditions prior to the start of the test. The neurobehavioral assay consisted of three phases: a 3-minute light phase (white light), a 3-minute dark phase (no light) and a 5 second UV phase (UV light only). The DanioVision system captured larval zebrafish movement throughout this assay at 30 frames per second.

### 2.8. Statistical analysis

LC50s for the LPS tests were determined using the Trimmed Spearmen-Karber method (Hamilton et al., 1977). The tail edema and mortality data from the initial and confirmatory drug testing was analyzed using GraphPad Prism 9 using Fisher’s Exact Test. Each drug treatment group was compared pairwise with the on-board positive control group on each test plate. Statistical significance was determined with values of *p* ≤ 0.05. Neurological behavioral testing data was analyzed by Ethovision software that produced an activity analysis profile per frame for each well of the 96 well plates. Any dead embryos were removed from the data set, and the remaining data set was imported into Excel. The offset function in Excel was used to average the percentage of pixel change per 0.1 s for each plate. These averages were then used to calculate the area under the curve (AUC) for each concentration and each phase of the neurological behavioral tests. The behavioral data was then statistically analyzed by drug per phase using a 2 tailed t-test to determine statistical significance *p* ≤ 0.01.

## 3. Results

### 3.1. Model optimization

Of the three bacterial strains of LPS tested, *K. pneumoniae* LPS was the most toxic, with a 24-hour exposure 5 dpf zebrafish LC50 of <25 μg/mL and a 24-hour exposure 3 dpf zebrafish LC50 of 34.6 μg/mL (95% confidence limits could not be calculated). The second most toxic LPS tested was *P. aeruginosa* with a 24-hour exposure to 5 dpf zebrafish LC50 of 67.5 μg/mL (95% confidence limits 61.8 - 73.8 μg/mL) and a 24-hour exposure to 3 dpf zebrafish LC50 of 64.71 μg/mL (95% confidence limits of 63.0 - 66.5 μg/mL). *E. coli* LPS was over an order of magnitude less toxic to zebrafish with an LC50 of >1000 ug/mL for both 24-hour exposures to the 5 dpf and the 3 dpf zebrafish embryos. (Figure 1).

**Figure 1:**
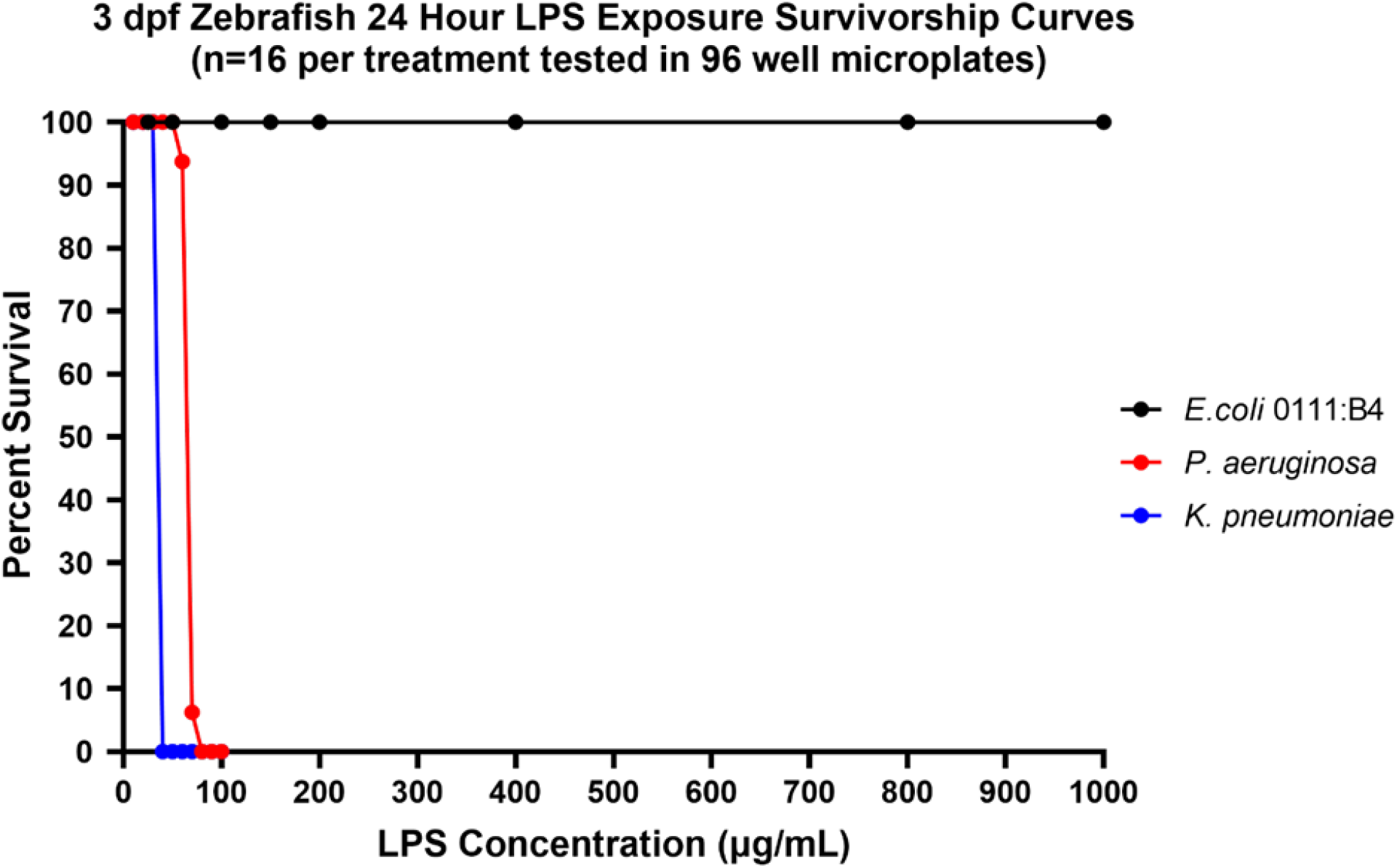
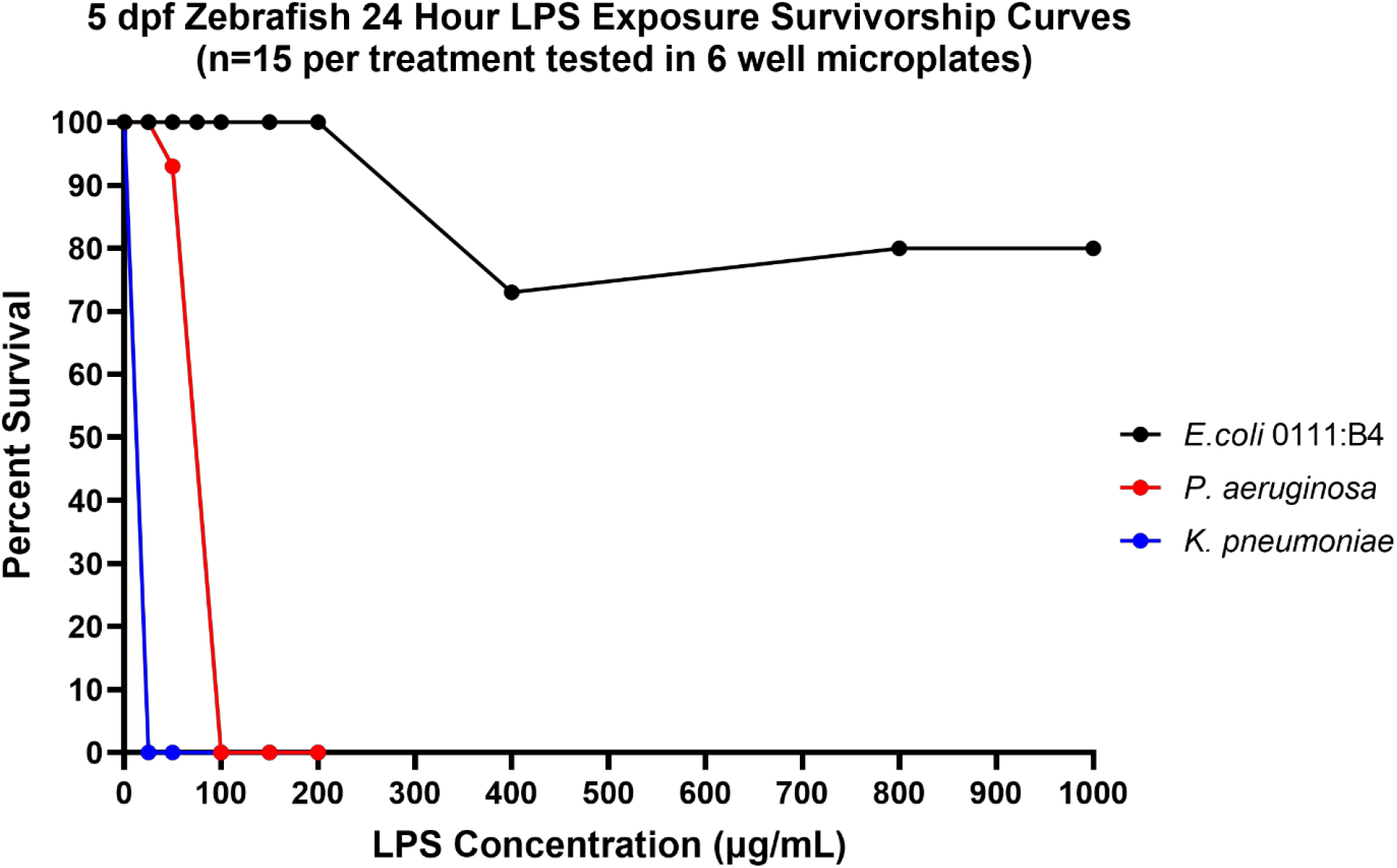
Dose response curves for 3 dpf zebrafish (top graph) and 5 dpf zebrafish (bottom graph) to 24 hour expsures to LPS derived from E.coli 0111:B4, P. aeruginosa, and K. pneumoniae.

*P. aeruginosa* LPS endpoint screening optimization was conducted to select a single LPS concentration that would produce a consistent endotoxic effect to facilitate reliable and repeatable drug compound screening studies. Specifically, the goal was to obtain near 50% mortality with 24-hour LPS exposed fish while generating a 100% combined response of tail edema and mortality. Figure 2 shows an example of a mild tail edema indictive of LPS endotoxicity in larval zebrafish. Figure 3 shows LPS dose response curves for tail edema and mortality during the protocol optimization and reference compound studies. Based on these response data, a concentration of 60 μg/mL *P. aeruginosa* LPS was found to be optimal for eliciting mortality and tail edema responses suitable for demonstrating the rescue potential of candidate drugs. While the ROS endpoint did show a quantifiable and statistically significant dose response to increasing concentrations of LPS (see supplemental data), it was not included as a drug screening endpoint. The additional time requirement for ROS measurements could not be justified given that mortality and tail edema together provided reliable and reproducible data sufficient for targeting efficacious drugs to prevent or treat sepsis.

**Figure 2:**
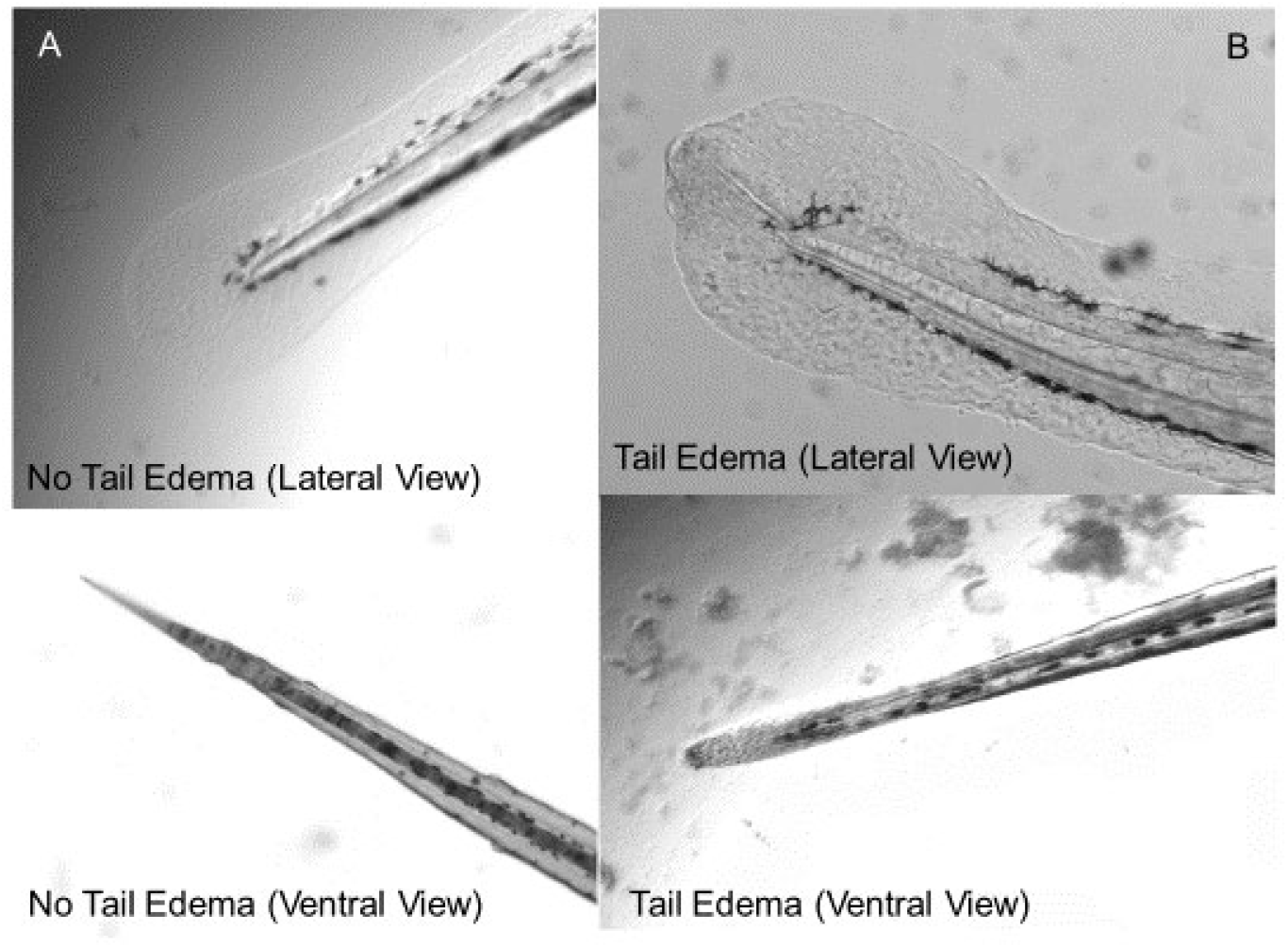
Caudal fin morphology of 3 dpf zebrafish. A. Images depict normal of caudal fin of control fish with no LPS exposure. B. Images depict the caudal fin after 24 hours of 60 μg/mL of *P. aeruginosa* LPS resulting in a tail emdema and vascular leakage.

**Figure 3:**
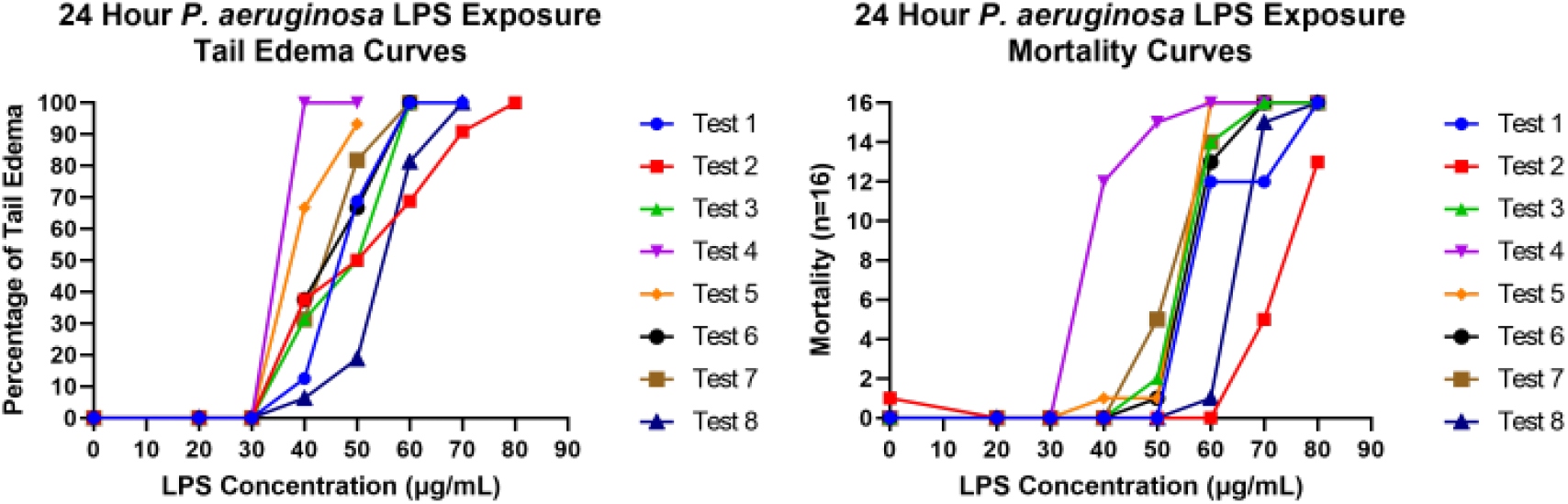
LPS dose response curves for replicate tests during the optimization phase of assay development. The left graph depicts the percentage of fish with tail edemas to increasing *P. aeruginosa* LPS concentrations. The right graph depicts of the number of dead fish to increasing *P. aeruginosa* LPS concentrations. The goal was to determine a concentration that yielded close to 50% mortality while eliciting 100% combined tail edema and mortality in all fish. A concentration of 60 μg/mL of P. aeruginosa LPS was chosen for follow-on chemical screening.

### 3.2. Model compound testing

Four model compounds (fasudil, hydrocortisone, dexamethasone, and fisetin) identified from prior published zebrafish LPS studies were evaluated. There was little change in LPS LC50 values with and without fisetin. LC50 values increased to 78.2 (95% confidence limits 72.6 – 84.3 μg/mL) from 73.6 μg/mL (95% confidence limits 68.7 – 78.8 μg/mL) with and without fisetin (400 μM) respectively and to 61.8 (95% confidence limits 57.4 – 66.5 μg/mL) from 60.5 μg/mL (95% confidence limits 56.3 – 65.0 μg/mL) with and without fasudil (10 μM). Neither compound showed complete rescue in any of the exposed fish at the 60 μg/mL LPS challenge concentration, although fasudil did show a statistically significant (p=0.0233) reduction in mortality at 24 hours post exposure. Prior work with fisetin by Molagoda et al. (2021) and other work by Hsu et al. (2018) used LPS microinjection methods and mortality rates in LPS positive controls for those studies were 30% (after 36 hours) and 5% (after 24 hours) respectively. The LPS challenge used in this study led to >60% mortality in LPS positive controls and thus direct comparisons are not advised given the differing exposure route and increased endotoxic challenge. Likewise, for fasudil, our selection of LPS (*P. aeruginosa* rather than *E. coli)* differed from prior work and may have been responsible for some of the differences in efficacy when compared to our study. For dexamethasone, the preliminary 6-hour post LPS-exposure reading demonstrated a statistically significant (p = 0.0091) reduction in tail edema and mortality occurred compared to the LPS control at 60 μg/mL, however, at 24-hours post LPS-exposure greater mortality with dexamethasone than without and there was no longer statically significant rescue from tail edema or mortality. The LPS 24-hour LC50 values were 48.2 μg/mL (95% confidence limits 45.75 – 50.68 μg/mL) with dexamethasone (100 μM) and 54.5 μg/mL (95% confidence limits 52.21 – 56.96 μg/mL) without dexamethasone. Hydrocortisone showed a similar response, with marginal mortality rescue noted at 6 hours post LPS exposure but with increased mortality over no treatment at 24 hours post LPS exposure (see supplemental data). No statistically significant difference was observed between LPS-exposed embryos treated with hydrocortisone and without.

### 3.3. Initial drug screening

Screening of the sepsis-focused custom FDA-approved drug library of 644 compounds yielded 84 compounds that had a statistically significant reduction in tail edema and mortality. To further down select top rescue drugs, a threshold of >60% rescue of tail edema and mortality was chosen. The number of compounds that met this threshold was 29. Statistical significance ranged from p = 0.0021 to < 0.0001 depending on the drug that had >60% rescue of tail edema and mortality. Of those 29 compounds, 4 compounds provided 100% rescue of tail edema and mortality (Ketanserin, Tegaserod, Brexpiprazole, and Dapoxetine). Figure 4 shows the response of Tegaserod along with other drugs from the initial screening library that were tested on the same 96 well plate. For this test plate, Tegaserod was the only drug that showed complete protection from LPS-induced tail edema and mortality (p-value < 0.0001). A summary of the full 644 compound screen efficacy results is provided in supplemental data.

**Figure 4:**
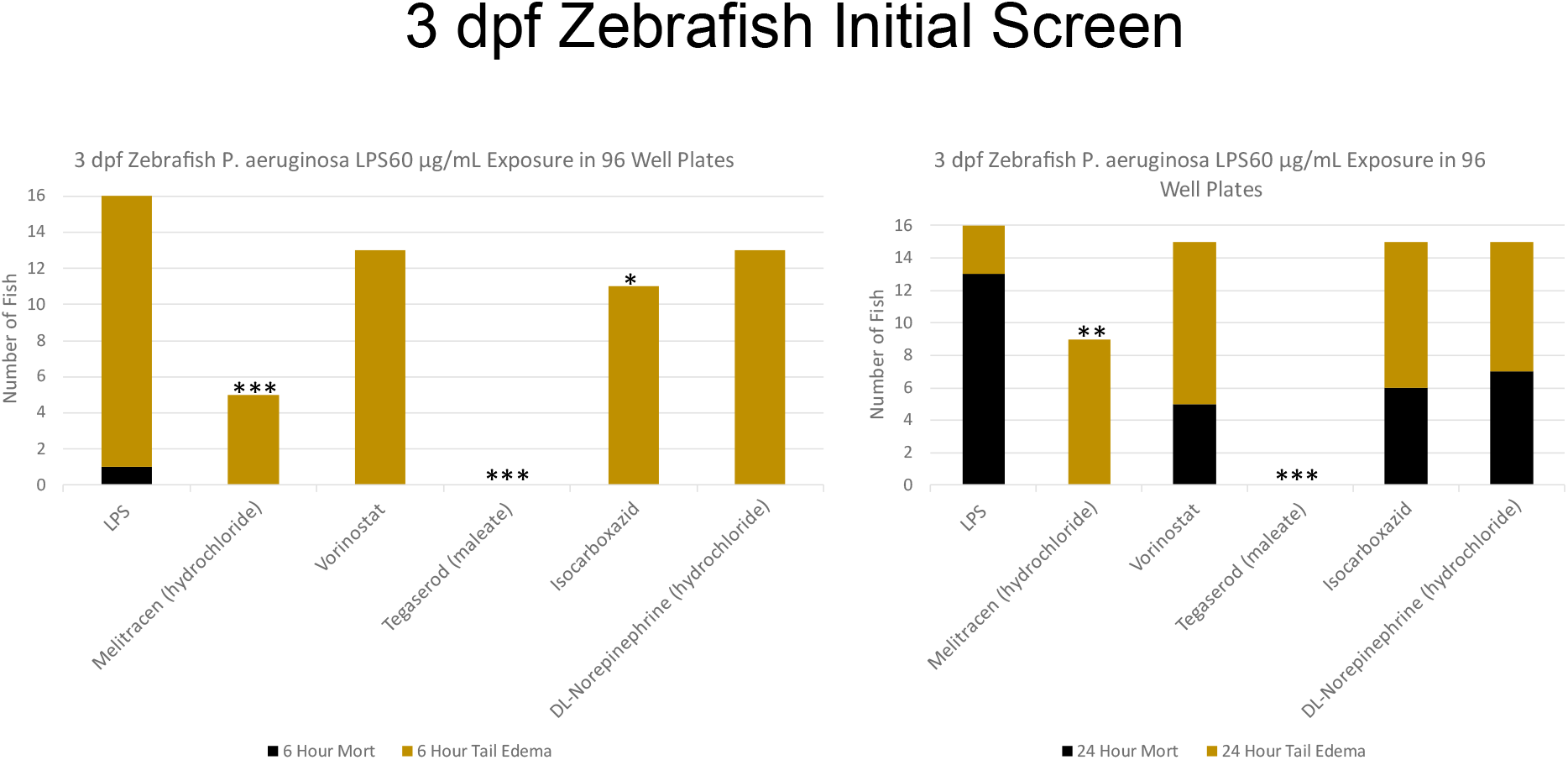
Example of the rescue results from the initial drug screenig from a single 96-well plate. Tegaserod showed complete resucue from LPS-induced tail and edema and mortality (n = 16 for each test condition) * = p-value ≤ 0.05, *\*\** = p-value ≤ 0.01, *** = p-value ≤ 0.001

Of the 29 compounds that exceeded 60% rescue, 19 came from the coagulation/anti-coagulation pathway library, 8 came from the mitochondria-targeted pathway library, and 2 each were from the cytokine targeted pathways and the highly selective targeted pathways. Two of the 29 top compounds were classified in more than one library. Sunitinib was in the mitochondria targeted library but also in the cytokine targeted library. Iproniazid was in the mitochondria target library as well as the highly selective library. Hit rates within a library were highest for the coagulation/anti-coagulation and the mitochondria-targeted pathway libraries as shown in table 1.

**Table 1.**

| Compound Library | Number of Compounds Evaluated | Number of Compounds Exceeding 60% Rescue | Hit Rate (>60% Rescue) |
| --- | --- | --- | --- |
| MedChemExpress Coagulation and Anti-Coagulation Targeted Pathways (FDA approved only) | 295 | 19 | 6.4 % |
| MedChemExpress Mitochondria-Targeted Pathways (FDA approved only) | 136 | 8 | 5.8 % |
| Selleckchem Cytokine Targeted Pathways (FDA approved only) | 140 | 2 | 1.4 % |
| Selleckchem Highly Selective Targeted Pathways (FDA approved only) | 114 | 2 | 1.7 % |

Note: Only 644 compounds were tested (not 685) as duplicates between pathway libraries were not tested.

### 3.4. Top rescue drug confirmatory testing and down selection

Confirmatory testing of the top 29 compounds showing >60% rescue in the initial drug screening effort yielded 16 compounds that demonstrated >80% rescue using 5 dpf larvae. Statistical significance for the compounds varied by compound from p-value = 0.0038 to < 0.0001. Five of the 16 compounds showed 100% rescue of tail edema and mortality at 5 dpf (Ketanserin tartrate, Tegaserod, Brexpiprazole, Sunitinib and Paliperidone palmitate). See supplemental data for a full summary.

Full drug efficacy dose range response curves for the top 16 compounds are shown in figure 5. Ketanserin and Tegaserod showed >60% rescue at drug concentrations as low as 0.01 μM, providing the widest therapeutic window when compared to other top rescue compounds, none of which had similar rescue below 1 μM.

**Figure 5:**
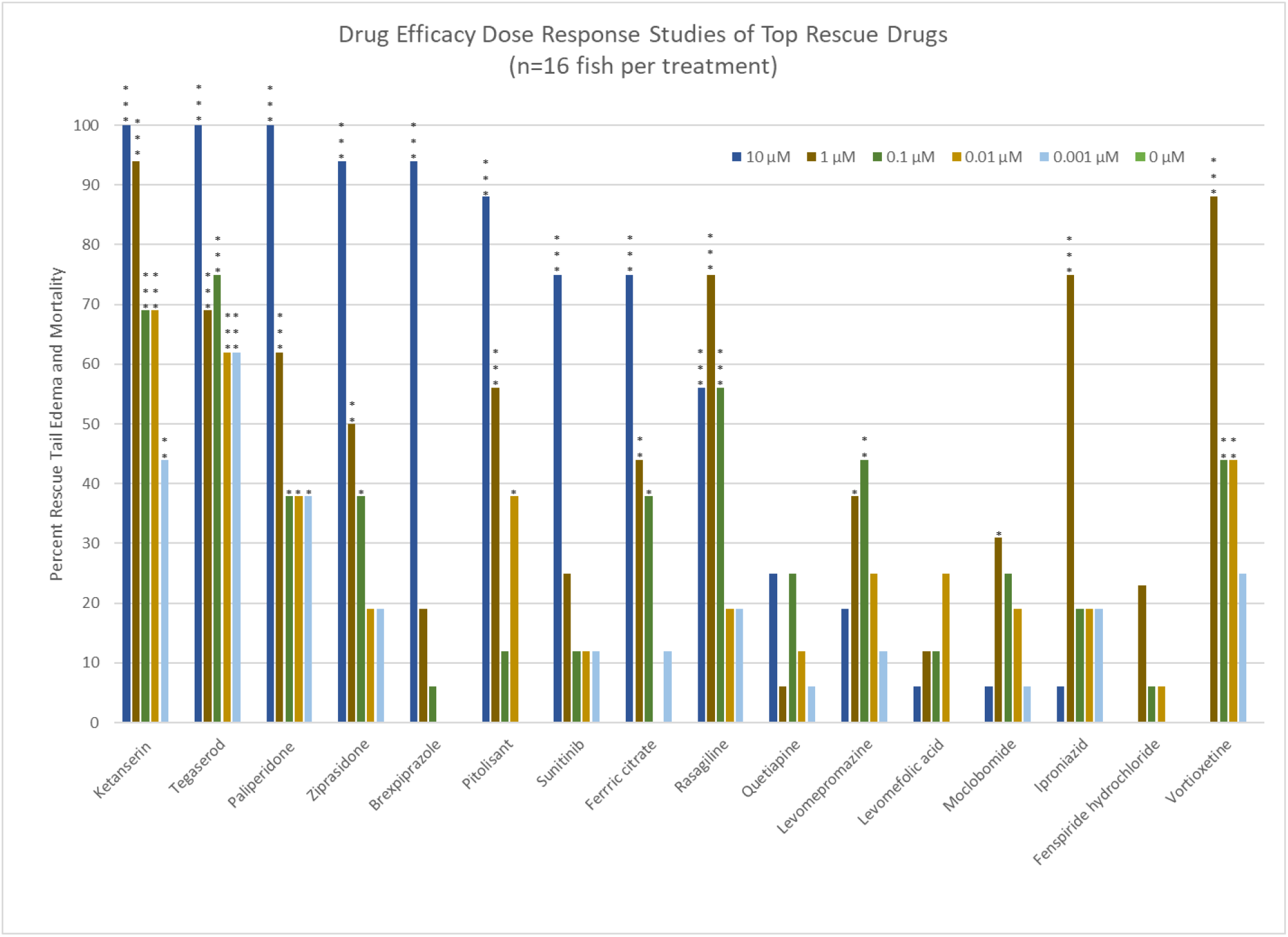
Drug efficacy dose range response curves for the top 16 rescue drugs. * = p-value ≤ 0.05, ** = p-value ≤ 0.01, and *** = p-value ≤ 0.001. n = 16 for each test condition.

Further evaluation of the top 16 rescue compounds used a 24-hour drug pre-treatment prior to 24-hour LPS exposure. Because of the increased LPS exposure in the pre-treatment dosing studies, there was increased LPS toxicity, leading to 100% mortality in the LPS positive controls. Drug pre-treatment generally didn’t improve rescue for most of the top drugs, other than Sunitinib and Ziprasidone which appeared to show improvement in rescue but were not statistically significant (Table 2). While drug pre-treatment generally did not improve rescue it also did not alter the efficacy and provides some early evidence that prophylactic use might be possible. The light/dark neurobehavioral testing (see supplemental data) identified 3 drugs (Ketanserin, Tegaserod, and Brexpiprazole) that showed little to no behavioral differences from negative controls. No significant differences were noted in any of these three drugs other than Brexpiprazole, which showed hyperactivity (p=0.005) during the dark phase with LPS exposure without 24-hour drug pre-treatment but was not significant when given prophylactically 24 hours prior to LPS exposure. Ketanserin, Tegaserod, and Brexpiprazole were further challenged with a lethal concentration of *K. pneumoniae* LPS (45 μg/mL). Both Brexpiprazole and Tegaserod showed 100% rescue of tail edema and mortality, while Ketanserin showed 88% rescue. Thus, these drugs show strong efficacy against endotoxicity not only from *P. aeruginosa* LPS, but also from *K. pneumoniae* LPS.

**Table 2.**
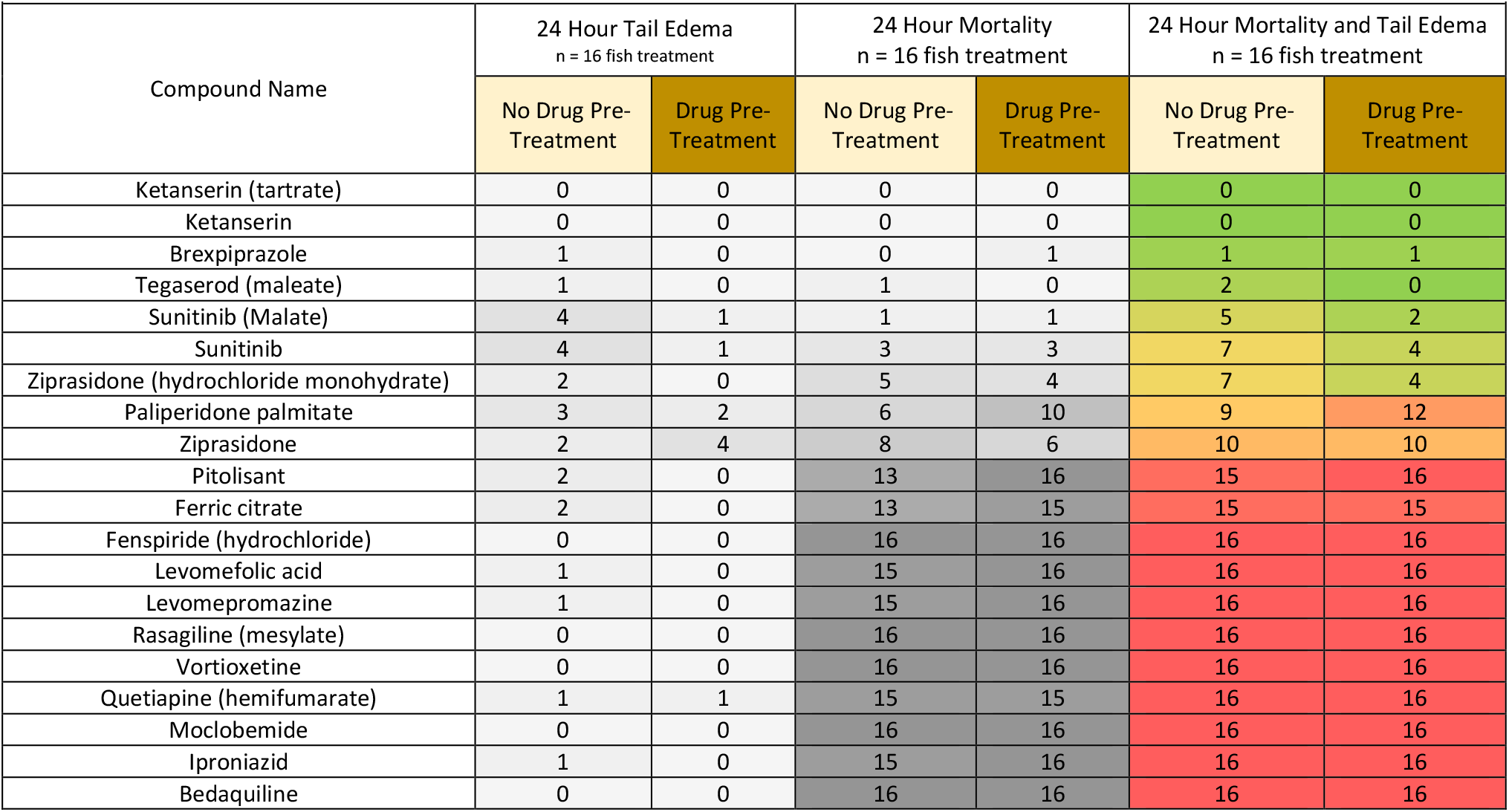

| Table 2 |  |  |  |  |  |  |
| --- | --- | --- | --- | --- | --- | --- |
| Compound Name | 24 Hour Tail Edema<br>n = 16 fish treatment |  | 24 Hour Mortality<br>n = 16 fish treatment |  | 24 Hour Mortality and Tail Edema<br>n = 16 fish treatment |  |
|  | No Drug Pre-Treatment | Drug Pre-Treatment | No Drug Pre-Treatment | Drug Pre-Treatment | No Drug Pre-Treatment | Drug Pre-Treatment |
| Ketanserin (tartrate) | 0 | 0 | 0 | 0 | 0 | 0 |
| Ketanserin | 0 | 0 | 0 | 0 | 0 | 0 |
| Brexpiprazole | 1 | 0 | 0 | 1 | 1 | 1 |
| Tegaserod (maleate) | 1 | 0 | 1 | 0 | 2 | 0 |
| Sunitinib (Malate) | 4 | 1 | 1 | 1 | 5 | 2 |
| Sunitinib | 4 | 1 | 3 | 3 | 7 | 4 |
| Ziprasidone (hydrochloride monohydrate) | 2 | 0 | 5 | 4 | 7 | 4 |
| Paliperidone palmitate | 3 | 2 | 6 | 10 | 9 | 12 |
| Ziprasidone | 2 | 4 | 8 | 6 | 10 | 10 |
| Pitolisant | 2 | 0 | 13 | 16 | 15 | 16 |
| Ferric citrate | 2 | 0 | 13 | 15 | 15 | 15 |
| Fenspiride (hydrochloride) | 0 | 0 | 16 | 16 | 16 | 16 |
| Levomefolic acid | 1 | 0 | 15 | 16 | 16 | 16 |
| Levomepromazine | 1 | 0 | 15 | 16 | 16 | 16 |
| Rasagiline (mesylate) | 0 | 0 | 16 | 16 | 16 | 16 |
| Vortioxetine | 0 | 0 | 16 | 16 | 16 | 16 |
| Quetiapine (hemifumarate) | 1 | 1 | 15 | 15 | 16 | 16 |
| Moclobemide | 0 | 0 | 16 | 16 | 16 | 16 |
| Iproniazid | 1 | 0 | 15 | 16 | 16 | 16 |
| Bedaquiline | 0 | 0 | 16 | 16 | 16 | 16 |

## 4. Discussion

Much has been published highlighting the role of cytokines and inflammatory pathways in the initiation and progression of sepsis. Although it was anticipated that the cytokine targeted pathways would yield a high level of positive hits for candidate sepsis rescue drugs, these pathways had the lowest number of hits of the targeted pathways screened in this study. Given the lack of efficacy of known anti-inflammatory drugs evaluated, this may not be unexpected. Likewise, modification of multiple cytokines might be required to affect sepsis progression, or cytokine pathways may be downstream of the true molecular initiation events that manifest in sepsis, septic shock and death. Eight of the 16 compounds demonstrating >80% rescue with 5 dpf larvae are known to target neuromodulatory pathways. Serotonin (5-HT receptors) pathway activity, in particular involving 5-HT2a, was present as a potential target in all 8 drugs but some also had activity within adrenergic, dopaminergic, muscarinic, and histaminergic pathways. Three compounds are monoamine oxidase inhibitors and three other top hits targeted potassium voltage-gated channels, apoptosis regulation, and tyrosine kinase inhibition. Current drug indications for the top 16 identified sepsis rescue drugs include 5 antipsychotics, 4 antidepressants, and single drugs identified for Parkinson’s disease, irritable bowel syndrome, anti-cancer/vascular endothelial and platelet growth factors, narcolepsy, respiratory disease, folate deficiency and iron supplementation/phosphate binder.

Of the top five rescue drugs (Table 3), only Ketanserin has been previously evaluated in a clinical trial as a potential sepsis therapy. In a 10-person sepsis clinical trial, Ketanserin improved microcirculation, but it’s vasodilatory activity may have hindered its transition to the clinic for the treatment of sepsis (Vellinga et al., 2015). Ketanserin may still have utility in situations where preventive treatment can be initiated early, prior to sepsis-associated hypotension. Except for Sunitinib, all five top rescue drugs have serotonergic (5-HT) receptor activity, which may indicate a potential common pathway for sepsis treatment and/or prevention. While Tegaserod, Brexpiprazole and Ziprasidone also target 5-HT2a receptors (and others), these drugs appear to lack the hypotension liability (as observed with Ketanserin), and thus potentially offer new options for sepsis prevention and treatment.

**Table 3.**

| Table 3 |  |  |  |  |  |
| --- | --- | --- | --- | --- | --- |
|  | No Pre-Treatment Rescue (n=16) | Pre-Treatment Rescue (n=16) | Post Test Behavioral Assay | Known Receptor Targets | Current Drug Use Owners (Brand Name) |
| Ketanserin | 100% | 100% | Normal | 5-HT<sub>2a</sub> (inverse agonist) | Jansen (Sufrexal) |
| Tegaserod | 88% | 100% | Normal | 5-HT<sub>4</sub> (partial agonist)<br>5-HT<sub>2a,b</sub> and c (antagonist) | Alfasigma (Zelnorm™) |
| Brexpiprazole | 94% | 94% | Normal | 5-HT<sub>1a</sub> (partial agonist)<br>5-HT<sub>2a</sub> (antagonist)<br>Noradrenergic α<sub>1b</sub> (antagonist)<br>Noradrenergic α<sub>2c</sub> (antagonist)<br>Dopamine D<sub>2</sub> (partial agonist) | Otsuka America (Rexulti®) |
| Sunitinib | 69% | 88% | Impaired neurobehavior from incomplete rescue of endotoxicity | Inhibits multiple receptor tyrosine kinases (RTKs) Platelet-derived growth factor receptor alpha and beta, Vascular endothelial growth factor receptor 1, 2 and 3, Mast/stem cell growth factor Kit, RTKe FLT3, Macrophage colony stimulating factor 1, Hepatocyte growth factor receptor | Pfizer (Sutent) |
| Ziprasidone | 56% | 75% | Impaired neurobehavior from incomplete rescue of endotoxicity | Dopamine D<sub>2</sub> (antagonist)<br>5-HT<sub>2a, c</sub> and d (antagonist)<br>5-HT<sub>1a</sub> (agonist) | Pfizer (Zeldox®) |

In addition to its well-known role within the central nervous system (CNS), 5-HT is of critical importance throughout the body, with complex modulatory roles in immune function and cellular signaling (Ahern, 2011, Zhang et al., 2017, Guan et al., 2022, Huang et al., 2022). 5-HT receptors also play a critical role in the cardiovascular system and in the regulation of endothelial barrier function and signaling; disruption of these functions has been implicated in the progression of sepsis (Ramage et. al 2008, Tanaka et. al 2021, Neumann et. al 2023). Further, prior work using a cecal ligation and puncture sepsis mouse model demonstrated that 5-HT proficient mice had a significantly lower survival rate than 5-HT-deficient mice (Zhang et al 2017), indicating that 5-HT pathways are likely involved in sepsis mortality. The known 5-HT activity of the discovered top rescue drugs, coupled with their ability to reduce LPS-induced vascular damage and mortality in the zebrafish endotoxemia model of sepsis, may indicate an ability to mediate endothelial function, which may be critical for silencing the signaling process involved in the progression of sepsis.

Of the 644 drugs in the screening library, a total of 103 drugs have known 5-HT receptor activity. Of these, only Ketanserin and Granisetron have been previously evaluated in clinical trials for sepsis. The Granisetron trial failed to improve 28-day mortality over placebo in a 150-person sepsis patient study (Guan et. al 2022). Granisetron is a 5HT3 receptor antagonist which differs in receptor activity from top rescue drugs discovered through the zebrafish endotoxemia sepsis model screening efforts. Granisetron showed protection against polymicrobial sepsis-induced acute lung injury in mice (Wang et al 2019) and sepsis-induced liver damage in rats (Aboyoussef et al., 2021). In zebrafish, Granisetron showed a significant reduction in mortality but did not rescue from vascular damage (tail edema) and thus was not selected as a leading sepsis drug candidate for further development. Recruitment for a sepsis clinical trial is currently underway for Ondansetron, another 5HT3 receptor antagonist included in the drug screening library but which was not efficacious in the zebrafish endotoxicity sepsis model.

Some results from this zebrafish sepsis model drug screening effort can be compared with findings from murine clinical studies. Published mouse LPS-induced sepsis studies showed that ciprofloxacin provided significant rescue from mortality (Ogino et al. 2009). In the zebrafish endotoxemia model used here, ciprofloxacin also provided rescue from mortality, but did not elicit rescue from vascular damage. Clinical trials for sepsis using anti-inflammatory drugs to date have not been successful (Ulloa et al., 2009). Similarly, corticosteroids such as dexamethasone, methylprednisolone, and hydrocortisone showed rescue early in the zebrafish sepsis model (6-hour exposure time point) but failed to show any level of rescue by 24 hours post exposure.

Zebrafish provide a reproducible model to evaluate sepsis therapy candidates with the additional benefit of a higher throughput over mammalian models. Neuromodulatory drugs may be viable targets for sepsis therapy. Candidates targeting 5-HT receptors have modulatory roles in the CNS but can also affect the cardiovascular system. Other identified complementary and novel targets also merit further study as sepsis therapeutics. The leading drug candidates identified in this study exceeded the therapeutic potential of commonly used compounds such as anti-inflammatory drugs and warrant follow-on evaluations using mammalian models of sepsis.

## CRediT authorship contribution statement

**Mark Widder:** Conceptualization, Methodology, Validation, Formal Analysis, Investigation, Resources, Data Curation, Writing – Original Draft, Visualization, Supervision, Project Administration, Funding Acquisition. **Chance Carbaugh:** Conceptualization, Methodology, Validation, Formal Analysis, Investigation, Resources, Data Curation, Software, Writing – Original Draft, Visualization. **William van der Schalie:** Conceptualization, Methodology, Data Curation, Writing – Original Draft, Visualization, Funding Acquisition. **Ronald Miller:** Investigation. **Linda Brennan:** Investigation. **Ashley Moore:** Validation, Investigation, Data Curation, Writing-review and editing. **Robert Campbell:** Methodology, Writing-review and editing. **Kevin Akers:** Conceptualization, Methodology. **Roseanne Ressner:** Methodology, Writing-review and editing. **Monica Martin:** Resources, Supervision. **Michael Madejczyk:** Resources, Supervision. **Blair Dancy:** Resources, Supervision. **Patricia Lee:** Resources, Supervision, Project Administration, Funding Acquisition. **Charlotte Lanteri:** Conceptualization, Methodology, Supervision, Project Administration, Funding Acquisition, Writing-review and editing.

## Acknowledgements

Material has been reviewed by the Walter Reed Army Institute of Research. There is no objection to its presentation and/or publication. The opinions or assertions contained herein are the private views of the authors, and are not to be construed as official, or as reflecting true views of the Department of the Army or the Department of Defense. Research was conducted under an IACUC-approved animal use protocol in an AAALAC International - accredited facility with a Public Health Services Animal Welfare Assurance and in compliance with the Animal Welfare Act and other federal statutes and regulations relating to laboratory animals.

