## Supplemental Data for "Identification of potential sepsis therapeutic drugs using a zebrafish rapid screening approach"

### Slide 1
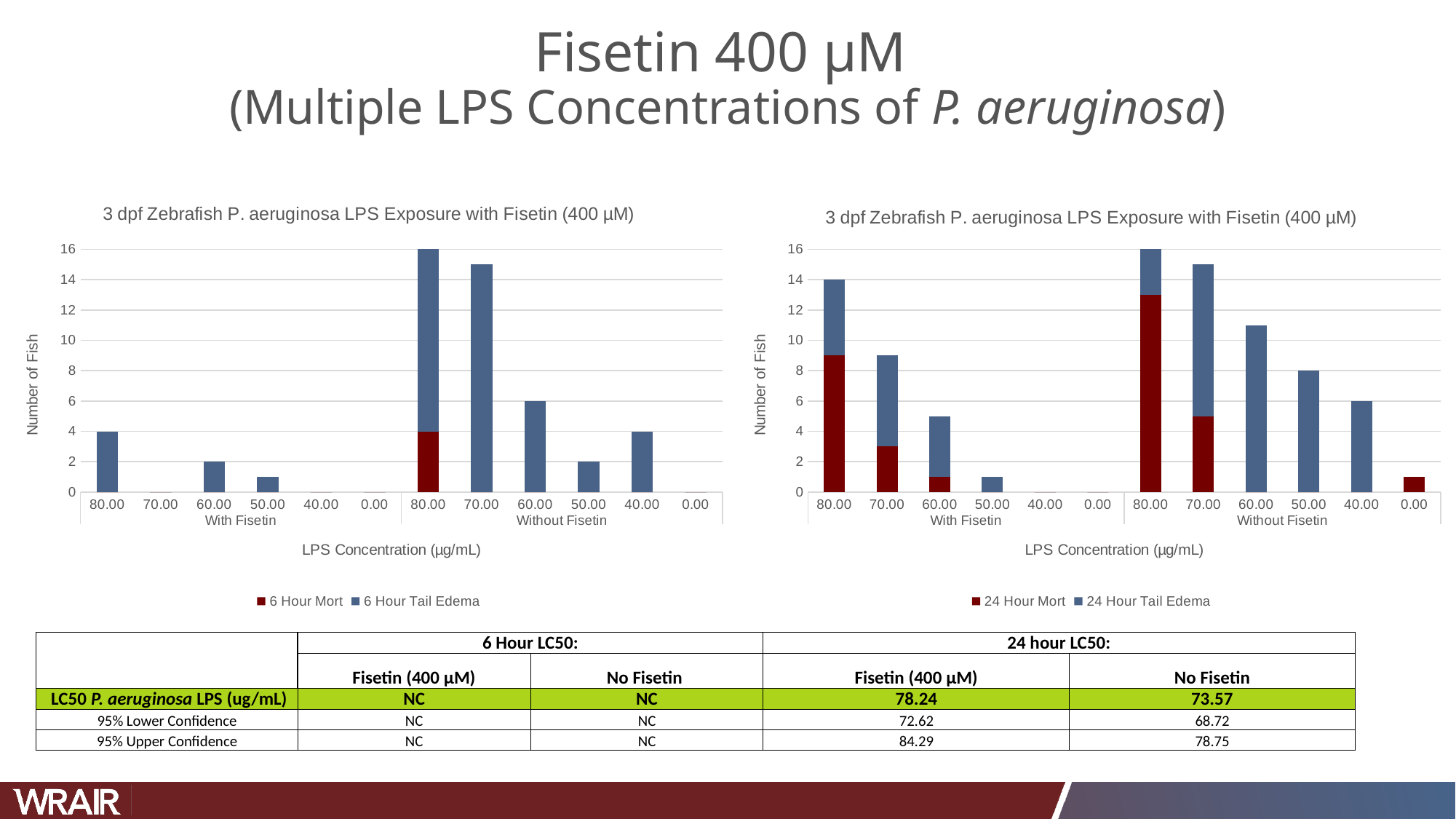

#### Fisetin 400 µM (Multiple LPS Concentrations of P. aeruginosa)
###### Chart:
| Category | 6 Hour Mort | 6 Hour Tail Edema |
|---|---|---|
| 80.00 | 0.0 | 4.0 |
| 70.00 | 0.0 | 0.0 |
| 60.00 | 0.0 | 2.0 |
| 50.00 | 0.0 | 1.0 |
| 40.00 | 0.0 | 0.0 |
| 0.00 | 0.0 | 0.0 |
| 80.00 | 4.0 | 12.0 |
| 70.00 | 0.0 | 15.0 |
| 60.00 | 0.0 | 6.0 |
| 50.00 | 0.0 | 2.0 |
| 40.00 | 0.0 | 4.0 |
| 0.00 | 0.0 | 0.0 |
###### Chart:
| Category | 24 Hour Mort | 24 Hour Tail Edema |
|---|---|---|
| 80.00 | 9.0 | 5.0 |
| 70.00 | 3.0 | 6.0 |
| 60.00 | 1.0 | 4.0 |
| 50.00 | 0.0 | 1.0 |
| 40.00 | 0.0 | 0.0 |
| 0.00 | 0.0 | 0.0 |
| 80.00 | 13.0 | 3.0 |
| 70.00 | 5.0 | 10.0 |
| 60.00 | 0.0 | 11.0 |
| 50.00 | 0.0 | 8.0 |
| 40.00 | 0.0 | 6.0 |
| 0.00 | 1.0 | 0.0 || | 6 Hour LC50: | | 24 hour LC50: |
| --- | --- | --- | --- | --- |
| | Fisetin (400 µM) | No Fisetin | Fisetin (400 µM) | No Fisetin |
| LC50 P. aeruginosa LPS (ug/mL) | NC | NC | 78.24 | 73.57 |
| 95% Lower Confidence | NC | NC | 72.62 | 68.72 |
| 95% Upper Confidence | NC | NC | 84.29 | 78.75 |

### Slide 2
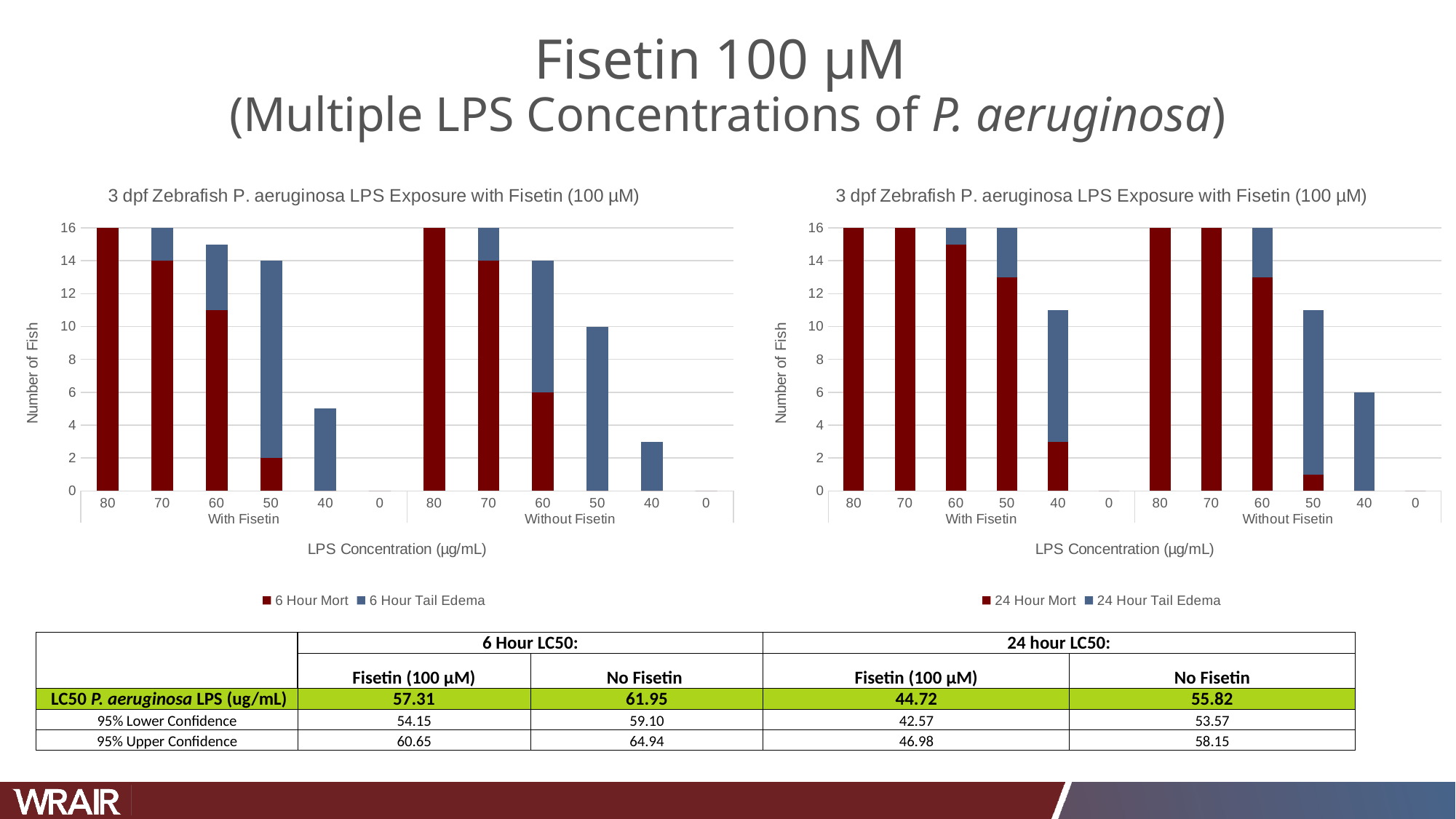

#### Fisetin 100 µM (Multiple LPS Concentrations of P. aeruginosa)
###### Chart:
| Category | 24 Hour Mort | 24 Hour Tail Edema |
|---|---|---|
| 80 | 16.0 | 0.0 |
| 70 | 16.0 | 0.0 |
| 60 | 15.0 | 1.0 |
| 50 | 13.0 | 3.0 |
| 40 | 3.0 | 8.0 |
| 0 | 0.0 | 0.0 |
| 80 | 16.0 | 0.0 |
| 70 | 16.0 | 0.0 |
| 60 | 13.0 | 3.0 |
| 50 | 1.0 | 10.0 |
| 40 | 0.0 | 6.0 |
| 0 | 0.0 | 0.0 |
###### Chart:
| Category | 6 Hour Mort | 6 Hour Tail Edema |
|---|---|---|
| 80 | 16.0 | 0.0 |
| 70 | 14.0 | 2.0 |
| 60 | 11.0 | 4.0 |
| 50 | 2.0 | 12.0 |
| 40 | 0.0 | 5.0 |
| 0 | 0.0 | 0.0 |
| 80 | 16.0 | 0.0 |
| 70 | 14.0 | 2.0 |
| 60 | 6.0 | 8.0 |
| 50 | 0.0 | 10.0 |
| 40 | 0.0 | 3.0 |
| 0 | 0.0 | 0.0 || | 6 Hour LC50: | | 24 hour LC50: |
| --- | --- | --- | --- | --- |
| | Fisetin (100 µM) | No Fisetin | Fisetin (100 µM) | No Fisetin |
| LC50 P. aeruginosa LPS (ug/mL) | 57.31 | 61.95 | 44.72 | 55.82 |
| 95% Lower Confidence | 54.15 | 59.10 | 42.57 | 53.57 |
| 95% Upper Confidence | 60.65 | 64.94 | 46.98 | 58.15 |

### Slide 3
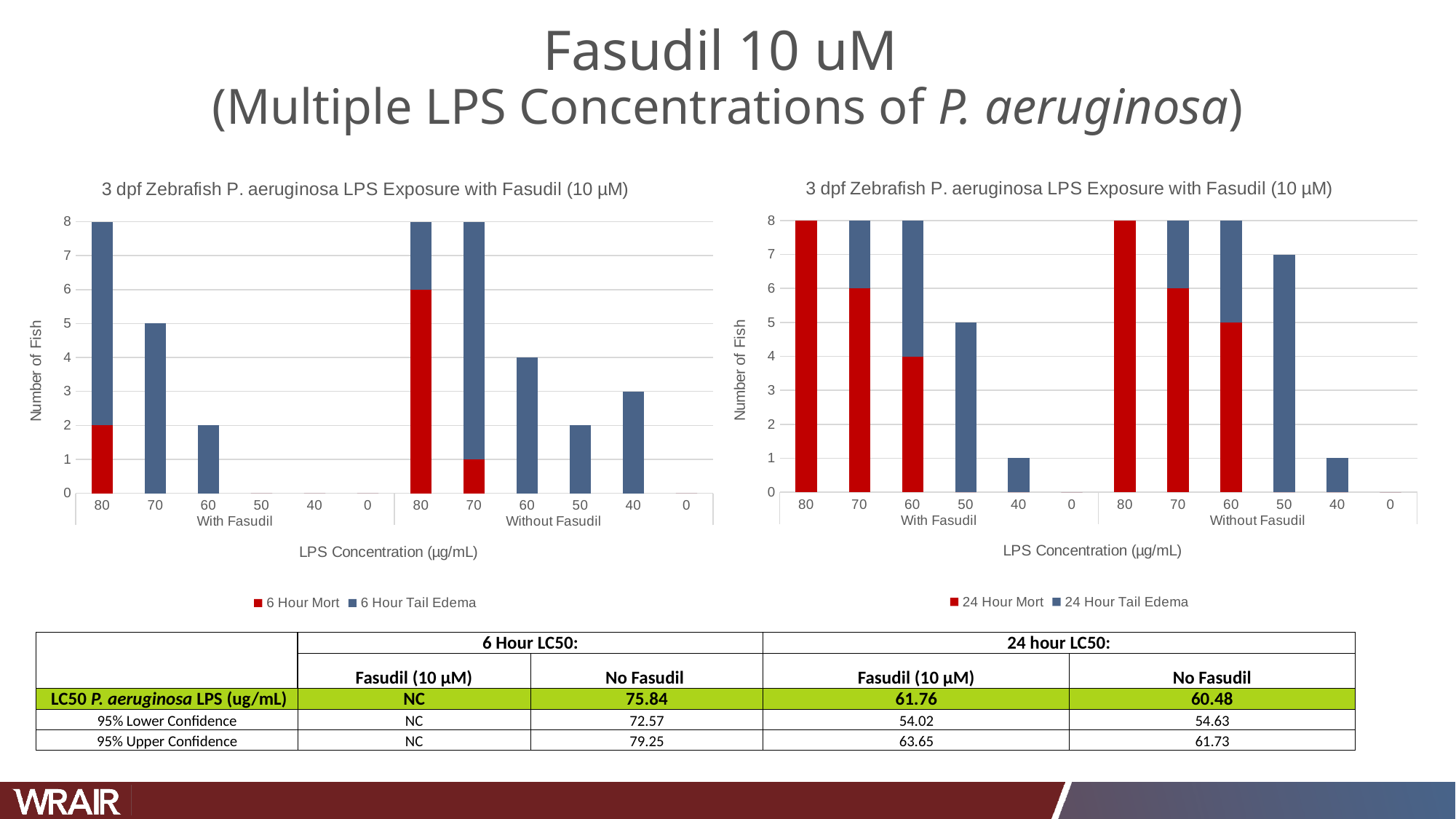

#### Fasudil 10 uM (Multiple LPS Concentrations of P. aeruginosa)
###### Chart:
| Category | 24 Hour Mort | 24 Hour Tail Edema |
|---|---|---|
| 80 | 8.0 | 0.0 |
| 70 | 6.0 | 2.0 |
| 60 | 4.0 | 4.0 |
| 50 | 0.0 | 5.0 |
| 40 | 0.0 | 1.0 |
| 0 | 0.0 | 0.0 |
| 80 | 8.0 | 0.0 |
| 70 | 6.0 | 2.0 |
| 60 | 5.0 | 3.0 |
| 50 | 0.0 | 7.0 |
| 40 | 0.0 | 1.0 |
| 0 | 0.0 | 0.0 |
###### Chart:
| Category | 6 Hour Mort | 6 Hour Tail Edema |
|---|---|---|
| 80 | 2.0 | 6.0 |
| 70 | 0.0 | 5.0 |
| 60 | 0.0 | 2.0 |
| 50 | 0.0 | 0.0 |
| 40 | 0.0 | 0.0 |
| 0 | 0.0 | 0.0 |
| 80 | 6.0 | 2.0 |
| 70 | 1.0 | 7.0 |
| 60 | 0.0 | 4.0 |
| 50 | 0.0 | 2.0 |
| 40 | 0.0 | 3.0 |
| 0 | 0.0 | 0.0 || | 6 Hour LC50: | | 24 hour LC50: |
| --- | --- | --- | --- | --- |
| | Fasudil (10 µM) | No Fasudil | Fasudil (10 µM) | No Fasudil |
| LC50 P. aeruginosa LPS (ug/mL) | NC | 75.84 | 61.76 | 60.48 |
| 95% Lower Confidence | NC | 72.57 | 54.02 | 54.63 |
| 95% Upper Confidence | NC | 79.25 | 63.65 | 61.73 |

### Slide 4
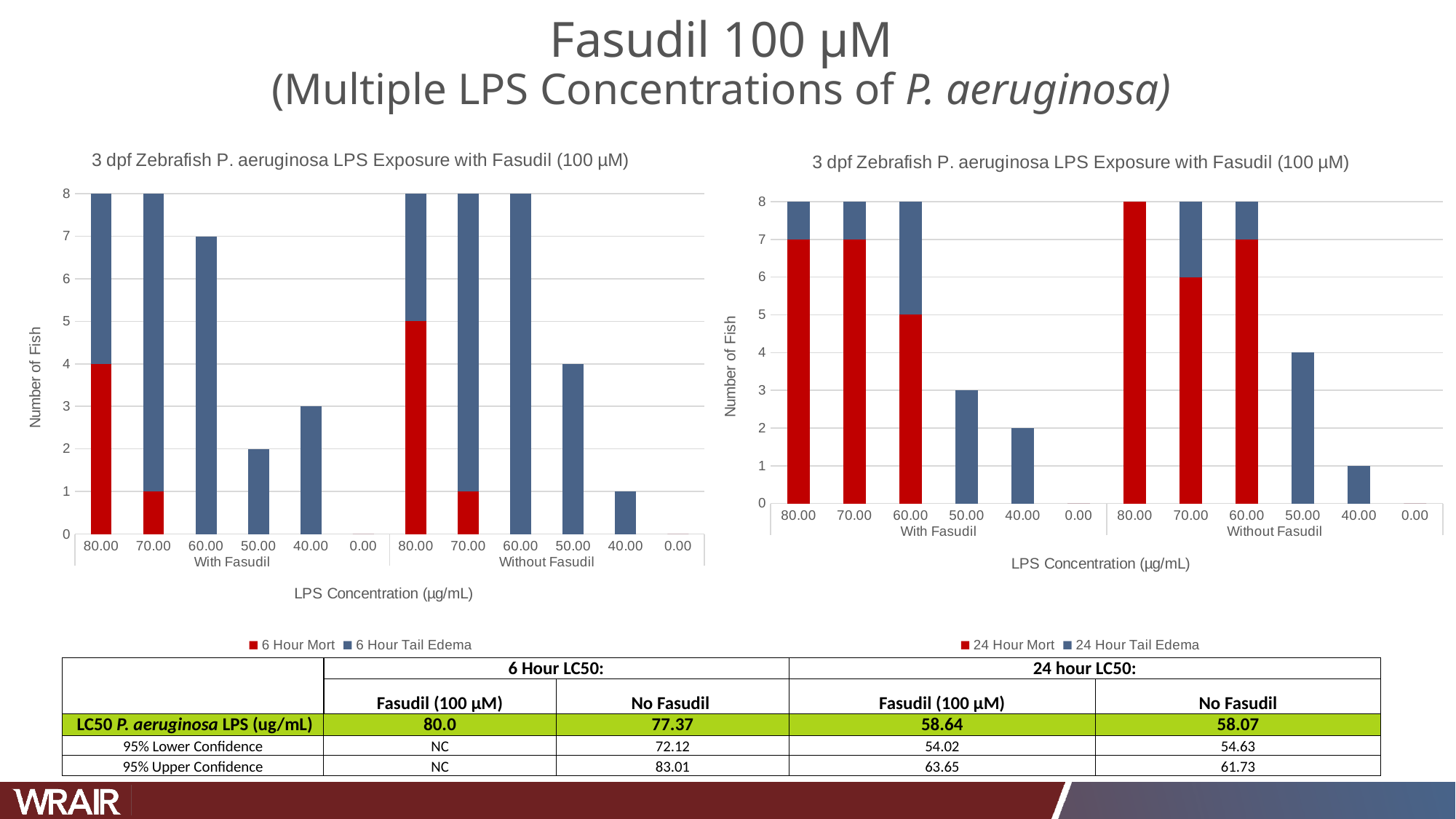

#### Fasudil 100 µM (Multiple LPS Concentrations of P. aeruginosa)
###### Chart:
| Category | 24 Hour Mort | 24 Hour Tail Edema |
|---|---|---|
| 80.00 | 7.0 | 1.0 |
| 70.00 | 7.0 | 1.0 |
| 60.00 | 5.0 | 3.0 |
| 50.00 | 0.0 | 3.0 |
| 40.00 | 0.0 | 2.0 |
| 0.00 | 0.0 | 0.0 |
| 80.00 | 8.0 | 0.0 |
| 70.00 | 6.0 | 2.0 |
| 60.00 | 7.0 | 1.0 |
| 50.00 | 0.0 | 4.0 |
| 40.00 | 0.0 | 1.0 |
| 0.00 | 0.0 | 0.0 |
###### Chart:
| Category | 6 Hour Mort | 6 Hour Tail Edema |
|---|---|---|
| 80.00 | 4.0 | 4.0 |
| 70.00 | 1.0 | 7.0 |
| 60.00 | 0.0 | 7.0 |
| 50.00 | 0.0 | 2.0 |
| 40.00 | 0.0 | 3.0 |
| 0.00 | 0.0 | 0.0 |
| 80.00 | 5.0 | 3.0 |
| 70.00 | 1.0 | 7.0 |
| 60.00 | 0.0 | 8.0 |
| 50.00 | 0.0 | 4.0 |
| 40.00 | 0.0 | 1.0 |
| 0.00 | 0.0 | 0.0 || | 6 Hour LC50: | | 24 hour LC50: |
| --- | --- | --- | --- | --- |
| | Fasudil (100 µM) | No Fasudil | Fasudil (100 µM) | No Fasudil |
| LC50 P. aeruginosa LPS (ug/mL) | 80.0 | 77.37 | 58.64 | 58.07 |
| 95% Lower Confidence | NC | 72.12 | 54.02 | 54.63 |
| 95% Upper Confidence | NC | 83.01 | 63.65 | 61.73 |

### Slide 5
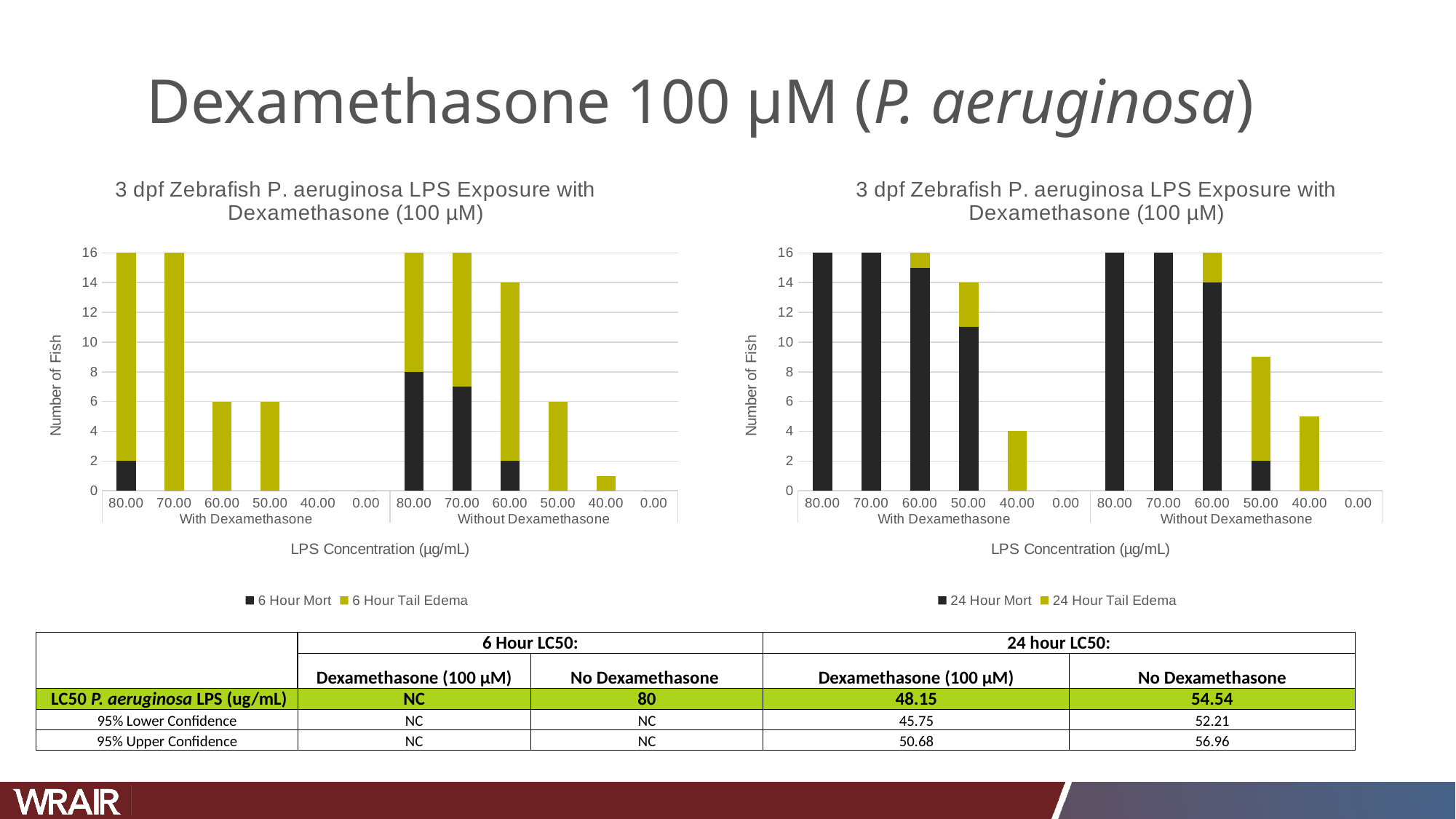

#### Dexamethasone 100 µM (P. aeruginosa)
###### Chart:
| Category | 6 Hour Mort | 6 Hour Tail Edema |
|---|---|---|
| 80.00 | 2.0 | 14.0 |
| 70.00 | 0.0 | 16.0 |
| 60.00 | 0.0 | 6.0 |
| 50.00 | 0.0 | 6.0 |
| 40.00 | 0.0 | 0.0 |
| 0.00 | 0.0 | 0.0 |
| 80.00 | 8.0 | 8.0 |
| 70.00 | 7.0 | 9.0 |
| 60.00 | 2.0 | 12.0 |
| 50.00 | 0.0 | 6.0 |
| 40.00 | 0.0 | 1.0 |
| 0.00 | 0.0 | 0.0 |
###### Chart:
| Category | 24 Hour Mort | 24 Hour Tail Edema |
|---|---|---|
| 80.00 | 16.0 | 0.0 |
| 70.00 | 16.0 | 0.0 |
| 60.00 | 15.0 | 1.0 |
| 50.00 | 11.0 | 3.0 |
| 40.00 | 0.0 | 4.0 |
| 0.00 | 0.0 | 0.0 |
| 80.00 | 16.0 | 0.0 |
| 70.00 | 16.0 | 0.0 |
| 60.00 | 14.0 | 2.0 |
| 50.00 | 2.0 | 7.0 |
| 40.00 | 0.0 | 5.0 |
| 0.00 | 0.0 | 0.0 || | 6 Hour LC50: | | 24 hour LC50: |
| --- | --- | --- | --- | --- |
| | Dexamethasone (100 µM) | No Dexamethasone | Dexamethasone (100 µM) | No Dexamethasone |
| LC50 P. aeruginosa LPS (ug/mL) | NC | 80 | 48.15 | 54.54 |
| 95% Lower Confidence | NC | NC | 45.75 | 52.21 |
| 95% Upper Confidence | NC | NC | 50.68 | 56.96 |

### Slide 6
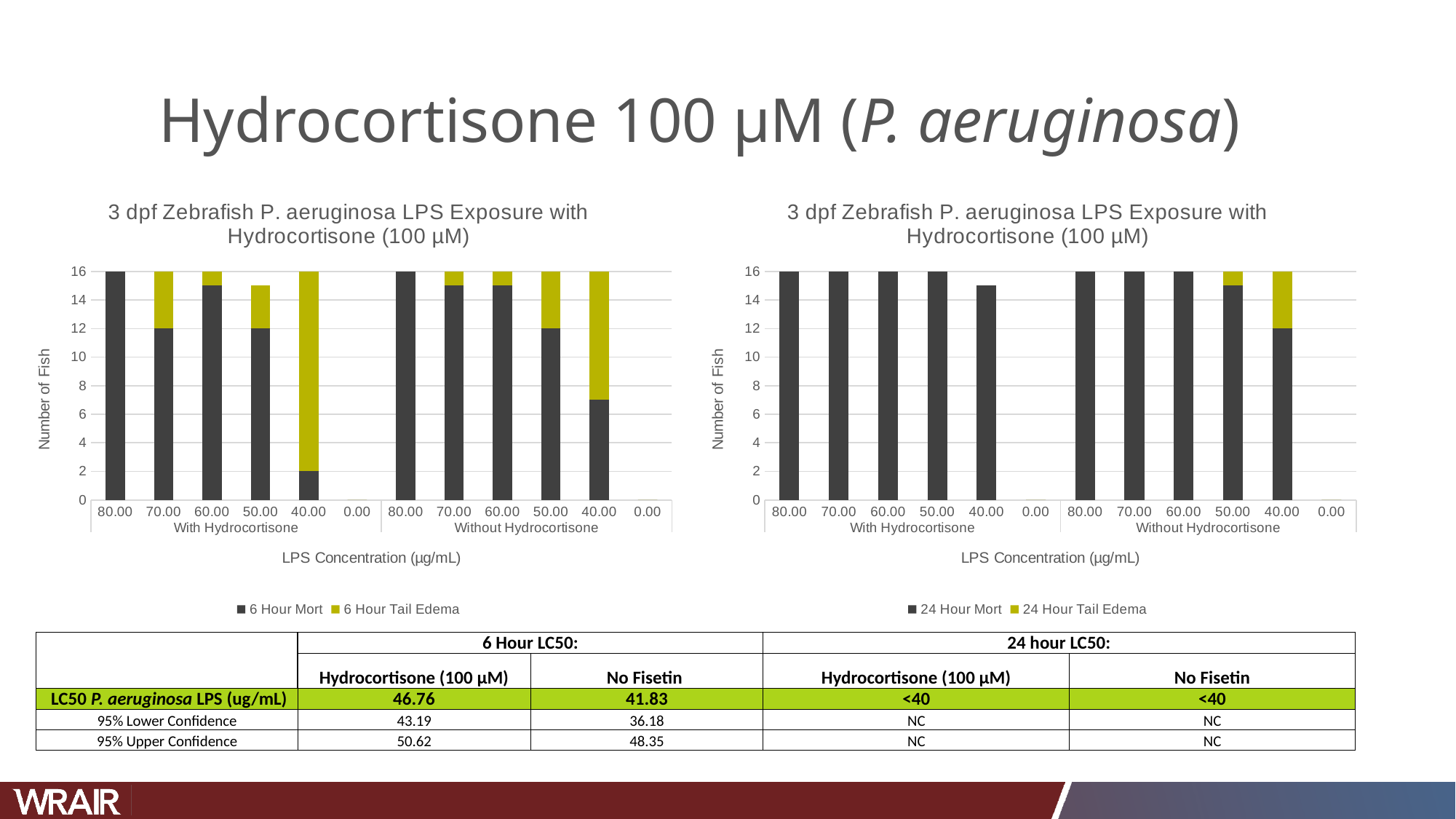

#### Hydrocortisone 100 µM (P. aeruginosa)
###### Chart:
| Category | 6 Hour Mort | 6 Hour Tail Edema |
|---|---|---|
| 80.00 | 16.0 | 0.0 |
| 70.00 | 12.0 | 4.0 |
| 60.00 | 15.0 | 1.0 |
| 50.00 | 12.0 | 3.0 |
| 40.00 | 2.0 | 14.0 |
| 0.00 | 0.0 | 0.0 |
| 80.00 | 16.0 | 0.0 |
| 70.00 | 15.0 | 1.0 |
| 60.00 | 15.0 | 1.0 |
| 50.00 | 12.0 | 4.0 |
| 40.00 | 7.0 | 9.0 |
| 0.00 | 0.0 | 0.0 |
###### Chart:
| Category | 24 Hour Mort | 24 Hour Tail Edema |
|---|---|---|
| 80.00 | 16.0 | 0.0 |
| 70.00 | 16.0 | 0.0 |
| 60.00 | 16.0 | 0.0 |
| 50.00 | 16.0 | 0.0 |
| 40.00 | 15.0 | 0.0 |
| 0.00 | 0.0 | 0.0 |
| 80.00 | 16.0 | 0.0 |
| 70.00 | 16.0 | 0.0 |
| 60.00 | 16.0 | 0.0 |
| 50.00 | 15.0 | 1.0 |
| 40.00 | 12.0 | 4.0 |
| 0.00 | 0.0 | 0.0 || | 6 Hour LC50: | | 24 hour LC50: |
| --- | --- | --- | --- | --- |
| | Hydrocortisone (100 µM) | No Fisetin | Hydrocortisone (100 µM) | No Fisetin |
| LC50 P. aeruginosa LPS (ug/mL) | 46.76 | 41.83 | <40 | <40 |
| 95% Lower Confidence | 43.19 | 36.18 | NC | NC |
| 95% Upper Confidence | 50.62 | 48.35 | NC | NC |

### Slide 7
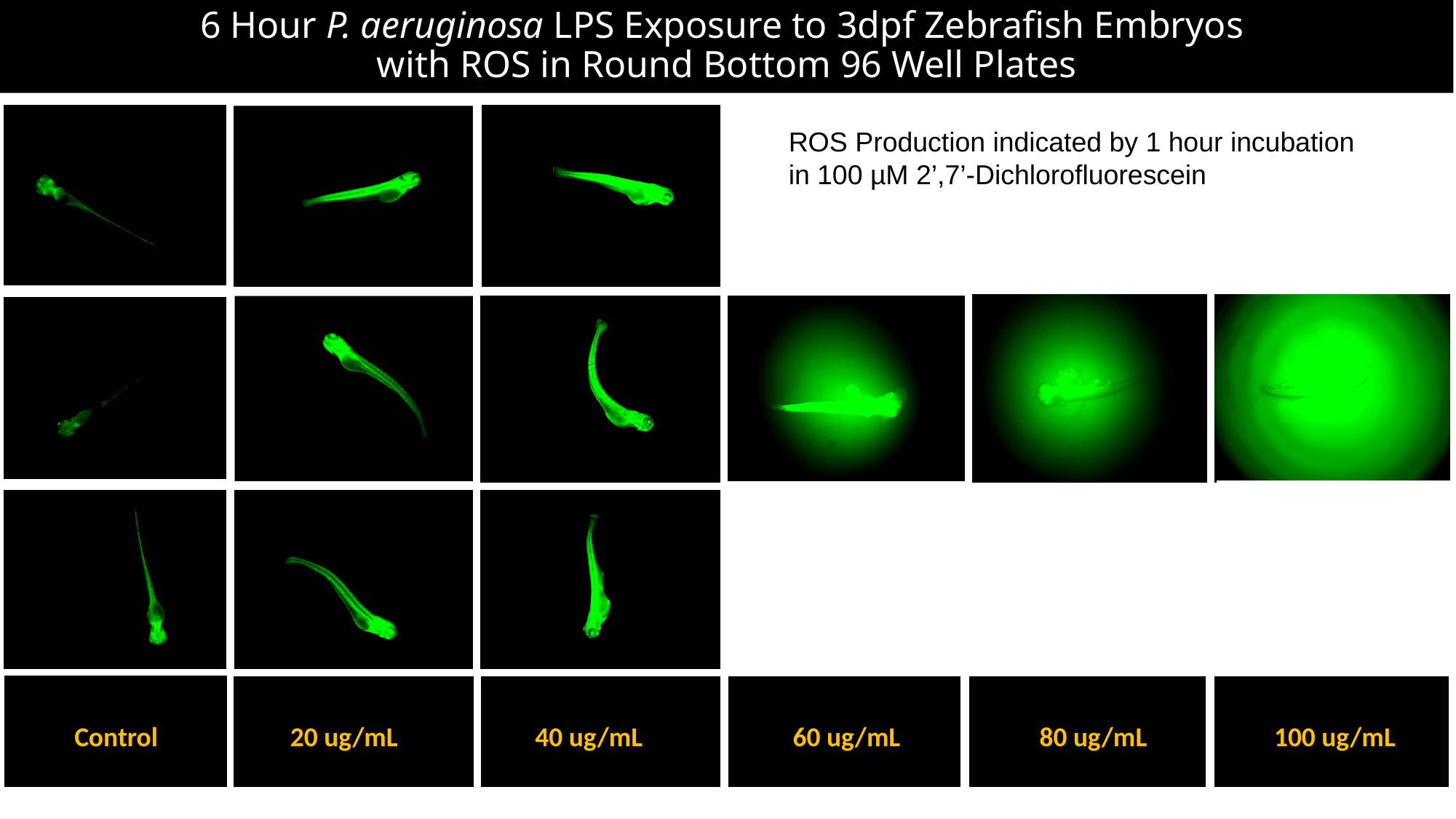

#### 6 Hour P. aeruginosa LPS Exposure to 3dpf Zebrafish Embryos with ROS in Round Bottom 96 Well Plates
ROS Production indicated by 1 hour incubation in 100 µM 2’,7’-Dichlorofluorescein
Control
20 ug/mL
40 ug/mL
60 ug/mL
80 ug/mL
100 ug/mL

### Slide 8
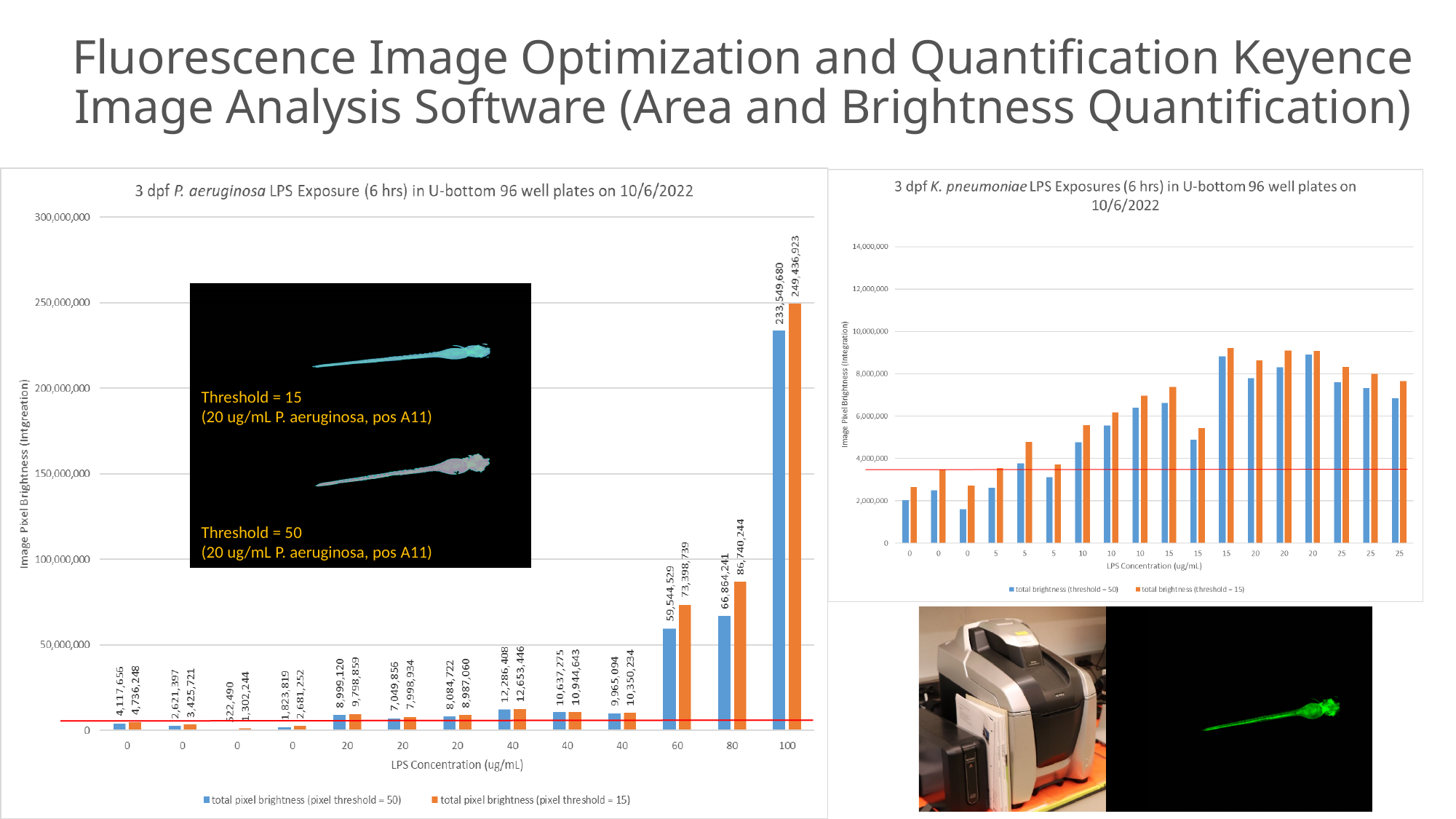

#### Fluorescence Image Optimization and Quantification Keyence Image Analysis Software (Area and Brightness Quantification)
Threshold = 15
(20 ug/mL P. aeruginosa, pos A11)
Threshold = 50
(20 ug/mL P. aeruginosa, pos A11)

### Slide 9
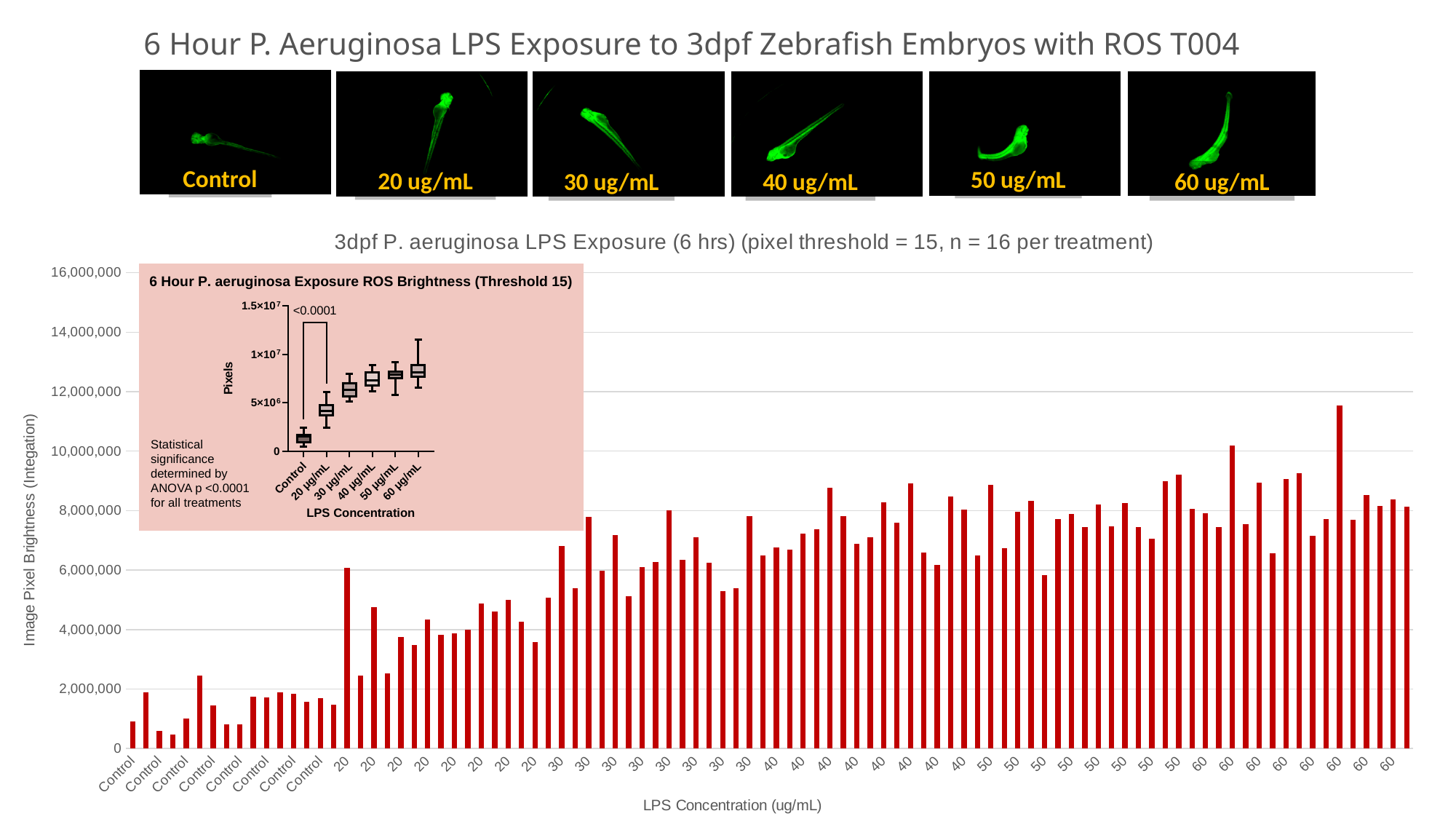

6 Hour P. Aeruginosa LPS Exposure to 3dpf Zebrafish Embryos with ROS T004
50 ug/mL
20 ug/mL
30 ug/mL
40 ug/mL
Control
60 ug/mL
###### Chart: 3dpf P. aeruginosa LPS Exposure (6 hrs) (pixel threshold = 15, n = 16 per treatment)
| Category | Brightness (integration) |
|---|---|
| Control | 910758.0 |
| Control | 1892653.0 |
| Control | 587714.0 |
| Control | 469017.0 |
| Control | 997075.0 |
| Control | 2445737.0 |
| Control | 1439374.0 |
| Control | 818807.0 |
| Control | 805457.0 |
| Control | 1733266.0 |
| Control | 1718534.0 |
| Control | 1885799.0 |
| Control | 1830727.0 |
| Control | 1563835.0 |
| Control | 1690960.0 |
| Control | 1477557.0 |
| 20 | 6076725.0 |
| 20 | 2444747.0 |
| 20 | 4749102.0 |
| 20 | 2523871.0 |
| 20 | 3759627.0 |
| 20 | 3478857.0 |
| 20 | 4340197.0 |
| 20 | 3827756.0 |
| 20 | 3875559.0 |
| 20 | 3996975.0 |
| 20 | 4885430.0 |
| 20 | 4605606.0 |
| 20 | 4988704.0 |
| 20 | 4255543.0 |
| 20 | 3574122.0 |
| 20 | 5084485.0 |
| 30 | 6822351.0 |
| 30 | 5385558.0 |
| 30 | 7785582.0 |
| 30 | 5989540.0 |
| 30 | 7173491.0 |
| 30 | 5133682.0 |
| 30 | 6098010.0 |
| 30 | 6277741.0 |
| 30 | 8019243.0 |
| 30 | 6353291.0 |
| 30 | 7111735.0 |
| 30 | 6244983.0 |
| 30 | 5291436.0 |
| 30 | 5401668.0 |
| 30 | 7810284.0 |
| 30 | 6490964.0 |
| 40 | 6767052.0 |
| 40 | 6684119.0 |
| 40 | 7231172.0 |
| 40 | 7365284.0 |
| 40 | 8777479.0 |
| 40 | 7817938.0 |
| 40 | 6889396.0 |
| 40 | 7102069.0 |
| 40 | 8280309.0 |
| 40 | 7605894.0 |
| 40 | 8903854.0 |
| 40 | 6599224.0 |
| 40 | 6166706.0 |
| 40 | 8475879.0 |
| 40 | 8028415.0 |
| 40 | 6494200.0 |
| 50 | 8877257.0 |
| 50 | 6736648.0 |
| 50 | 7953002.0 |
| 50 | 8334961.0 |
| 50 | 5829774.0 |
| 50 | 7716560.0 |
| 50 | 7880116.0 |
| 50 | 7441074.0 |
| 50 | 8207035.0 |
| 50 | 7465527.0 |
| 50 | 8253696.0 |
| 50 | 7442055.0 |
| 50 | 7055294.0 |
| 50 | 9000668.0 |
| 50 | 9214674.0 |
| 50 | 8059185.0 |
| 60 | 7921638.0 |
| 60 | 7435755.0 |
| 60 | 10176664.0 |
| 60 | 7549524.0 |
| 60 | 8943677.0 |
| 60 | 6566626.0 |
| 60 | 9059720.0 |
| 60 | 9267307.0 |
| 60 | 7151812.0 |
| 60 | 7706199.0 |
| 60 | 11532670.0 |
| 60 | 7683733.0 |
| 60 | 8519938.0 |
| 60 | 8162231.0 |
| 60 | 8374024.0 |
| 60 | 8140930.0 |Statistical significance determined by ANOVA p <0.0001 for all treatments

### Slide 10
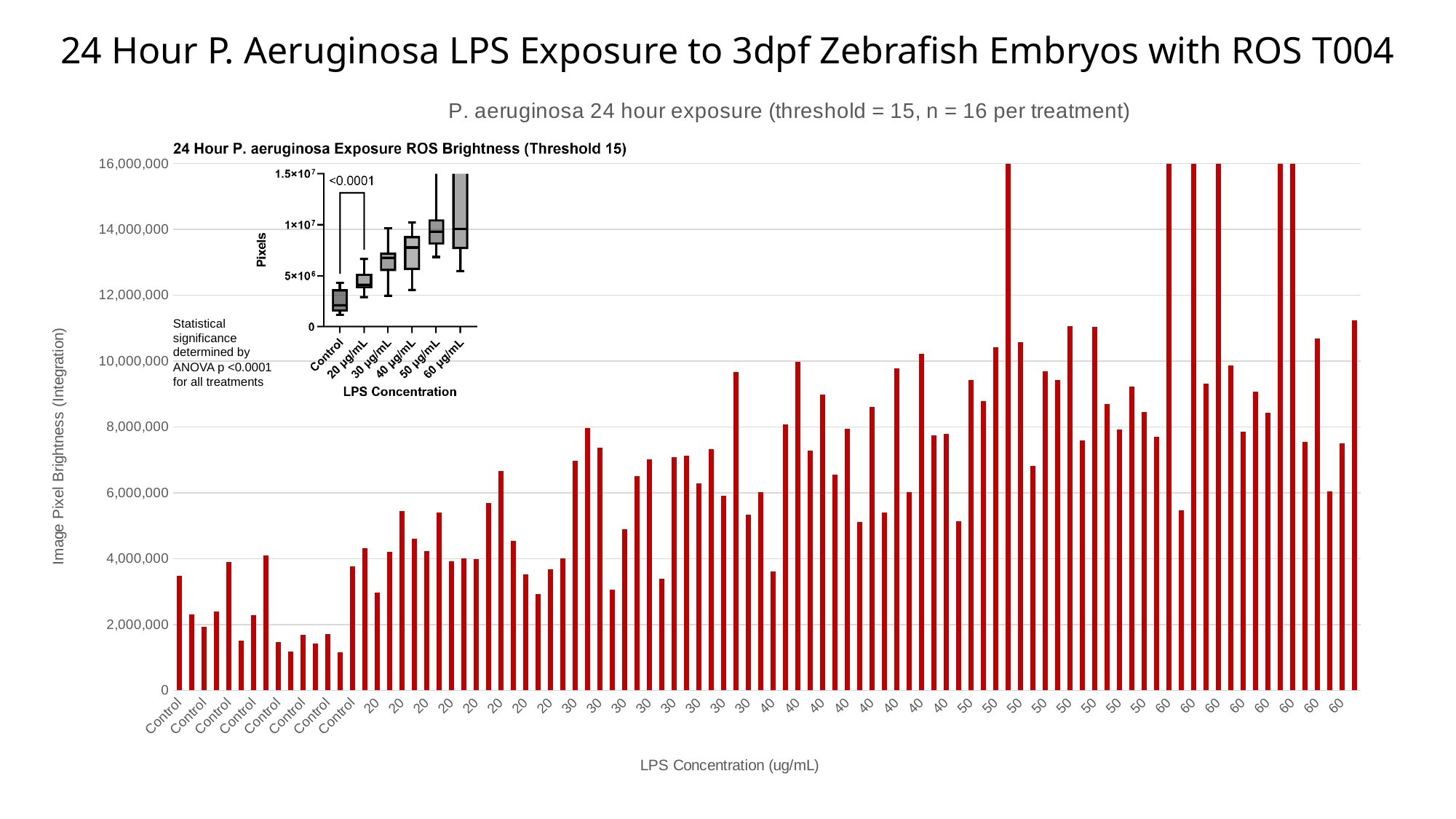

24 Hour P. Aeruginosa LPS Exposure to 3dpf Zebrafish Embryos with ROS T004
###### Chart: P. aeruginosa 24 hour exposure (threshold = 15, n = 16 per treatment)
| Category | Brightness (integration) |
|---|---|
| Control | 3488035.0 |
| Control | 2297140.0 |
| Control | 1933167.0 |
| Control | 2402138.0 |
| Control | 3904649.0 |
| Control | 1520968.0 |
| Control | 2282730.0 |
| Control | 4103210.0 |
| Control | 1461369.0 |
| Control | 1170126.0 |
| Control | 1681327.0 |
| Control | 1418656.0 |
| Control | 1711991.0 |
| Control | 1164365.0 |
| Control | 3767675.0 |
| Control | 4313118.0 |
| 20 | 2980622.0 |
| 20 | 4209063.0 |
| 20 | 5454153.0 |
| 20 | 4604723.0 |
| 20 | 4226593.0 |
| 20 | 5401119.0 |
| 20 | 3922352.0 |
| 20 | 4018058.0 |
| 20 | 3980137.0 |
| 20 | 5693621.0 |
| 20 | 6656594.0 |
| 20 | 4533962.0 |
| 20 | 3522326.0 |
| 20 | 2919890.0 |
| 20 | 3685002.0 |
| 20 | 4011994.0 |
| 30 | 6966334.0 |
| 30 | 7963975.0 |
| 30 | 7360493.0 |
| 30 | 3049159.0 |
| 30 | 4893016.0 |
| 30 | 6516260.0 |
| 30 | 7021377.0 |
| 30 | 3386434.0 |
| 30 | 7086723.0 |
| 30 | 7122626.0 |
| 30 | 6275975.0 |
| 30 | 7331213.0 |
| 30 | 5900128.0 |
| 30 | 9671979.0 |
| 30 | 5326381.0 |
| 30 | 6014717.0 |
| 40 | 3614899.0 |
| 40 | 8070564.0 |
| 40 | 9988913.0 |
| 40 | 7270558.0 |
| 40 | 8988658.0 |
| 40 | 6548988.0 |
| 40 | 7951843.0 |
| 40 | 5124582.0 |
| 40 | 8616142.0 |
| 40 | 5408824.0 |
| 40 | 9779215.0 |
| 40 | 6014917.0 |
| 40 | 10222642.0 |
| 40 | 7748926.0 |
| 40 | 7790896.0 |
| 40 | 5137501.0 |
| 50 | 9424217.0 |
| 50 | 8779425.0 |
| 50 | 10417149.0 |
| 50 | 187510925.0 |
| 50 | 10578191.0 |
| 50 | 6827309.0 |
| 50 | 9695327.0 |
| 50 | 9434309.0 |
| 50 | 11061100.0 |
| 50 | 7593150.0 |
| 50 | 11045650.0 |
| 50 | 8688850.0 |
| 50 | 7916060.0 |
| 50 | 9218220.0 |
| 50 | 8454717.0 |
| 50 | 7693061.0 |
| 60 | 295966838.0 |
| 60 | 5473230.0 |
| 60 | 130464143.0 |
| 60 | 9314437.0 |
| 60 | 267847304.0 |
| 60 | 9877064.0 |
| 60 | 7852238.0 |
| 60 | 9083629.0 |
| 60 | 8430290.0 |
| 60 | 294053761.0 |
| 60 | 300240876.0 |
| 60 | 7545095.0 |
| 60 | 10687269.0 |
| 60 | 6046478.0 |
| 60 | 7494586.0 |
| 60 | 11248348.0 |
Statistical significance determined by ANOVA p <0.0001 for all treatments

### Slide 11
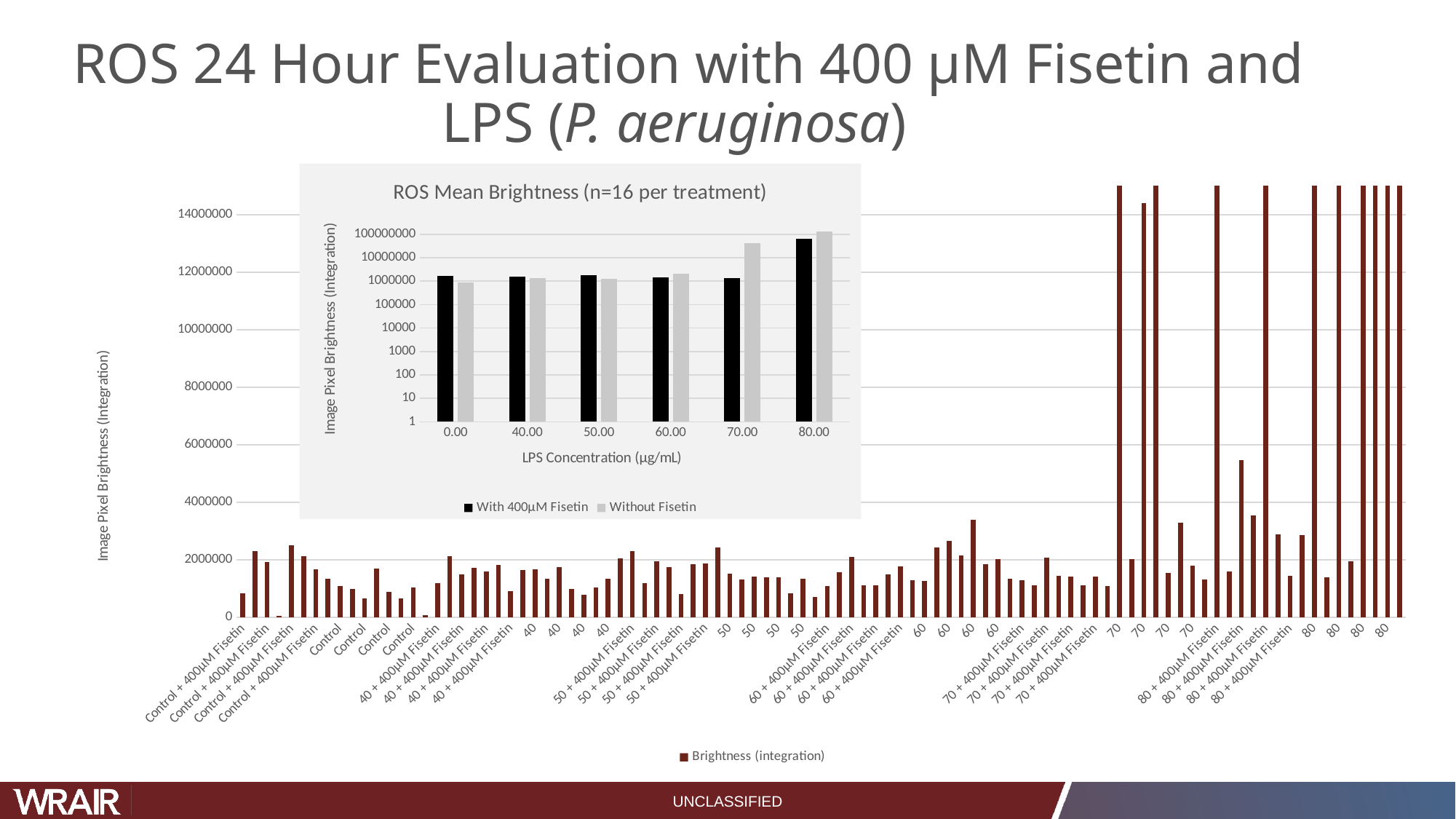

#### ROS 24 Hour Evaluation with 400 µM Fisetin and LPS (P. aeruginosa)
###### Chart: ROS Mean Brightness (n=16 per treatment)
| Category | With 400µM Fisetin | Without Fisetin |
|---|---|---|
| 0 | 1635921.2857142857 | 887799.875 |
| 40 | 1569854.0 | 1370966.0 |
| 50 | 1773981.625 | 1239551.375 |
| 60 | 1443051.125 | 2138479.25 |
| 70 | 1372059.5 | 42815977.25 |
| 80 | 62877304.25 | 130246858.125 |
###### Chart
| Category | Brightness (integration) |
|---|---|
| Control + 400µM Fisetin | 848416.0 |
| Control + 400µM Fisetin | 2316062.0 |
| Control + 400µM Fisetin | 1928673.0 |
| Control + 400µM Fisetin | 43906.0 |
| Control + 400µM Fisetin | 2505581.0 |
| Control + 400µM Fisetin | 2127777.0 |
| Control + 400µM Fisetin | 1681034.0 |
| Control + 400µM Fisetin | 1338646.0 |
| Control | 1089158.0 |
| Control | 982125.0 |
| Control | 656896.0 |
| Control | 1691837.0 |
| Control | 894273.0 |
| Control | 657016.0 |
| Control | 1043478.0 |
| Control | 87616.0 |
| 40 + 400µM Fisetin | 1203736.0 |
| 40 + 400µM Fisetin | 2137051.0 |
| 40 + 400µM Fisetin | 1502015.0 |
| 40 + 400µM Fisetin | 1735924.0 |
| 40 + 400µM Fisetin | 1602874.0 |
| 40 + 400µM Fisetin | 1821326.0 |
| 40 + 400µM Fisetin | 919247.0 |
| 40 + 400µM Fisetin | 1636659.0 |
| 40 | 1662805.0 |
| 40 | 1337470.0 |
| 40 | 1757496.0 |
| 40 | 992113.0 |
| 40 | 782364.0 |
| 40 | 1046018.0 |
| 40 | 1337549.0 |
| 40 | 2051913.0 |
| 50 + 400µM Fisetin | 2311507.0 |
| 50 + 400µM Fisetin | 1178820.0 |
| 50 + 400µM Fisetin | 1941406.0 |
| 50 + 400µM Fisetin | 1759361.0 |
| 50 + 400µM Fisetin | 822698.0 |
| 50 + 400µM Fisetin | 1861421.0 |
| 50 + 400µM Fisetin | 1873962.0 |
| 50 + 400µM Fisetin | 2442678.0 |
| 50 | 1519268.0 |
| 50 | 1318690.0 |
| 50 | 1415292.0 |
| 50 | 1383695.0 |
| 50 | 1390427.0 |
| 50 | 844191.0 |
| 50 | 1340785.0 |
| 50 | 704063.0 |
| 60 + 400µM Fisetin | 1085742.0 |
| 60 + 400µM Fisetin | 1561428.0 |
| 60 + 400µM Fisetin | 2100355.0 |
| 60 + 400µM Fisetin | 1116914.0 |
| 60 + 400µM Fisetin | 1118473.0 |
| 60 + 400µM Fisetin | 1488527.0 |
| 60 + 400µM Fisetin | 1774011.0 |
| 60 + 400µM Fisetin | 1298959.0 |
| 60 | 1258218.0 |
| 60 | 2427770.0 |
| 60 | 2670106.0 |
| 60 | 2151696.0 |
| 60 | 3387284.0 |
| 60 | 1843022.0 |
| 60 | 2015846.0 |
| 60 | 1353892.0 |
| 70 + 400µM Fisetin | 1288729.0 |
| 70 + 400µM Fisetin | 1117200.0 |
| 70 + 400µM Fisetin | 2089234.0 |
| 70 + 400µM Fisetin | 1437905.0 |
| 70 + 400µM Fisetin | 1418891.0 |
| 70 + 400µM Fisetin | 1121630.0 |
| 70 + 400µM Fisetin | 1423027.0 |
| 70 + 400µM Fisetin | 1079860.0 |
| 70 | 191654439.0 |
| 70 | 2036733.0 |
| 70 | 14410115.0 |
| 70 | 126468205.0 |
| 70 | 1539089.0 |
| 70 | 3282469.0 |
| 70 | 1808154.0 |
| 70 | 1328614.0 |
| 80 + 400µM Fisetin | 301872194.0 |
| 80 + 400µM Fisetin | 1593843.0 |
| 80 + 400µM Fisetin | 5460279.0 |
| 80 + 400µM Fisetin | 3553403.0 |
| 80 + 400µM Fisetin | 183363414.0 |
| 80 + 400µM Fisetin | 2880697.0 |
| 80 + 400µM Fisetin | 1445074.0 |
| 80 + 400µM Fisetin | 2849530.0 |
| 80 | 178267174.0 |
| 80 | 1385364.0 |
| 80 | 153708721.0 |
| 80 | 1940202.0 |
| 80 | 245133063.0 |
| 80 | 124349384.0 |
| 80 | 183179176.0 |
| 80 | 154011781.0 |UNCLASSIFIED

### Slide 12
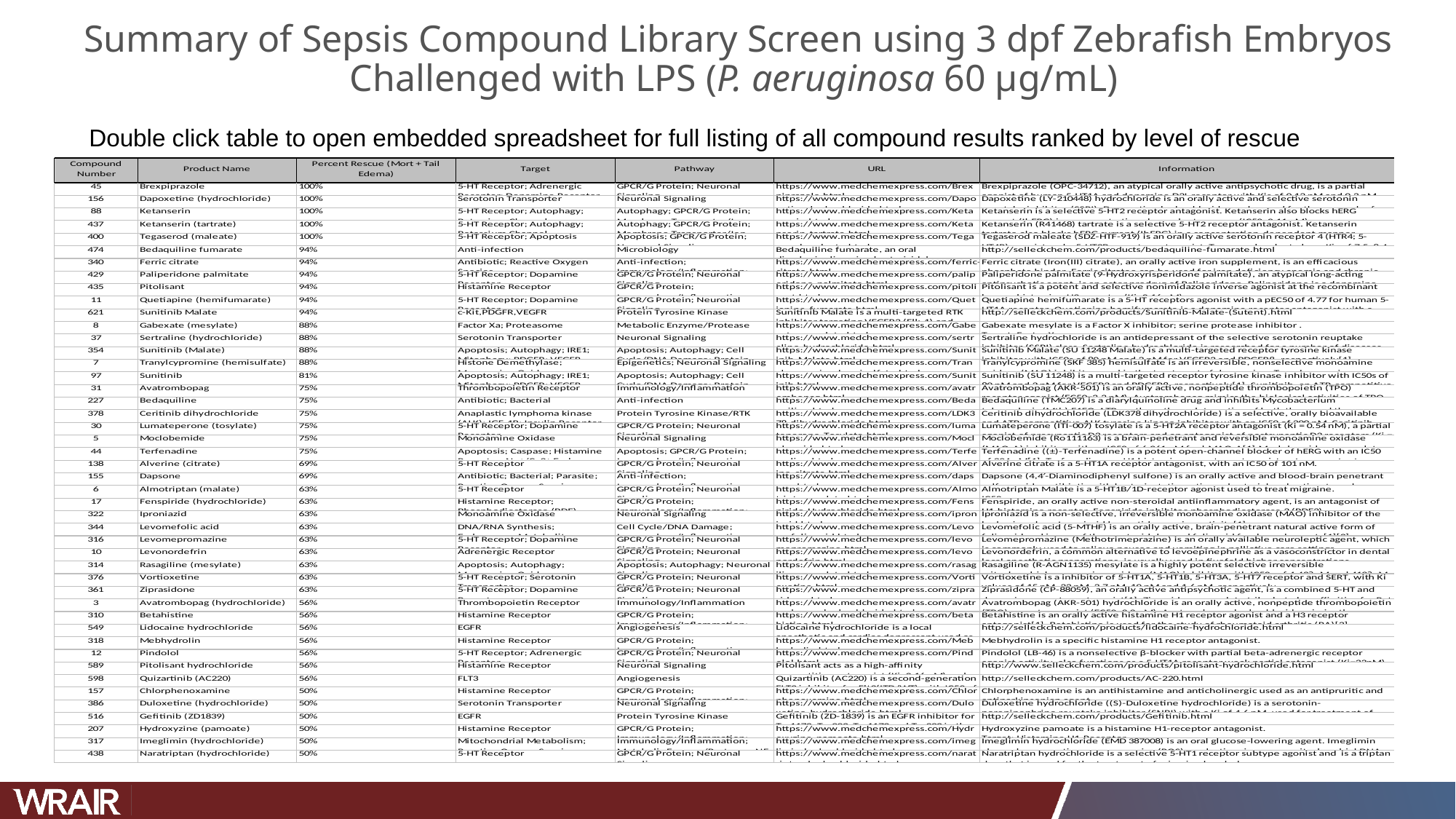

#### Summary of Sepsis Compound Library Screen using 3 dpf Zebrafish Embryos Challenged with LPS (P. aeruginosa 60 µg/mL)
Double click table to open embedded spreadsheet for full listing of all compound results ranked by level of rescue

### Slide 13
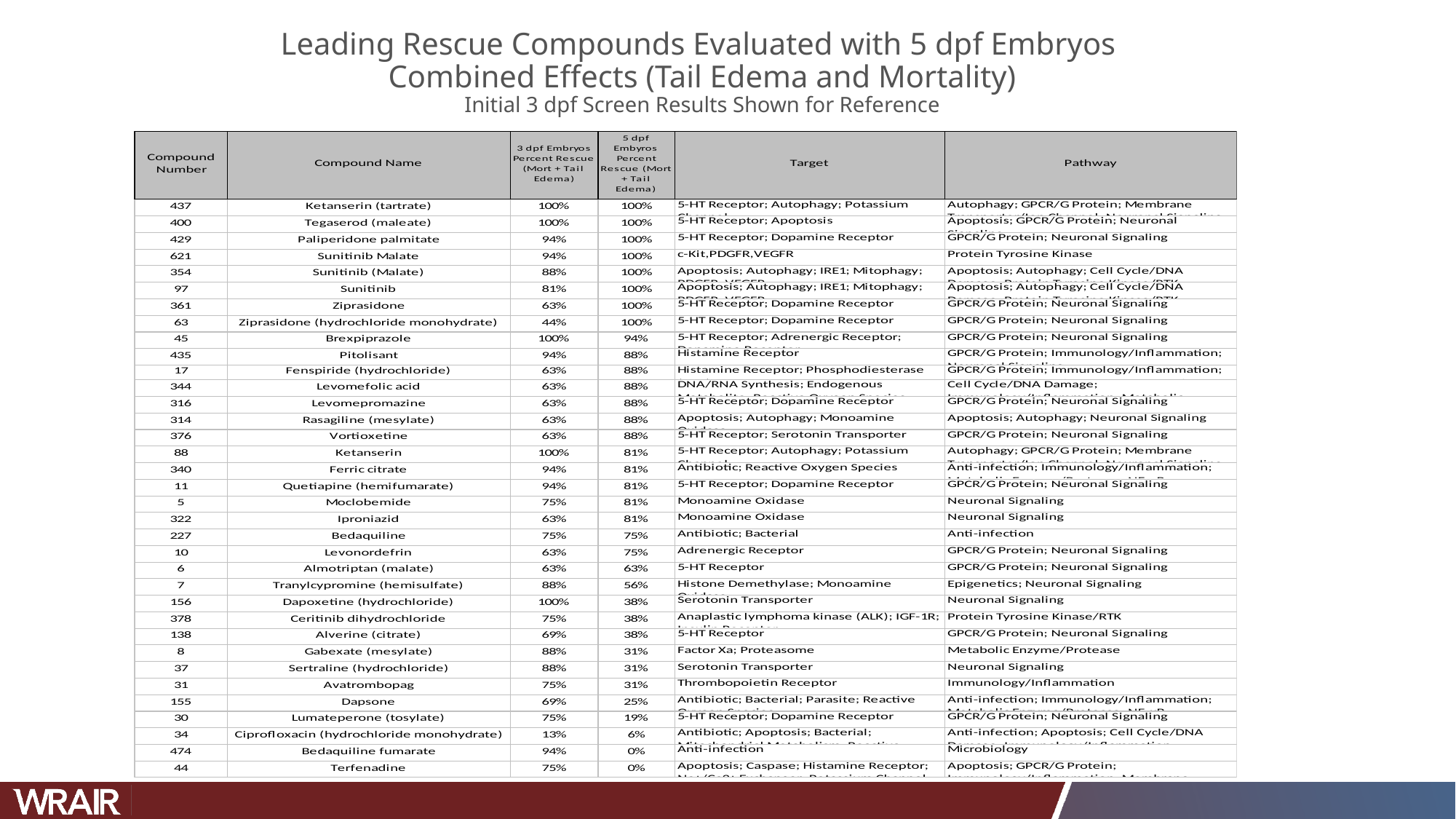

#### Leading Rescue Compounds Evaluated with 5 dpf Embryos Combined Effects (Tail Edema and Mortality) Initial 3 dpf Screen Results Shown for Reference

### Slide 14
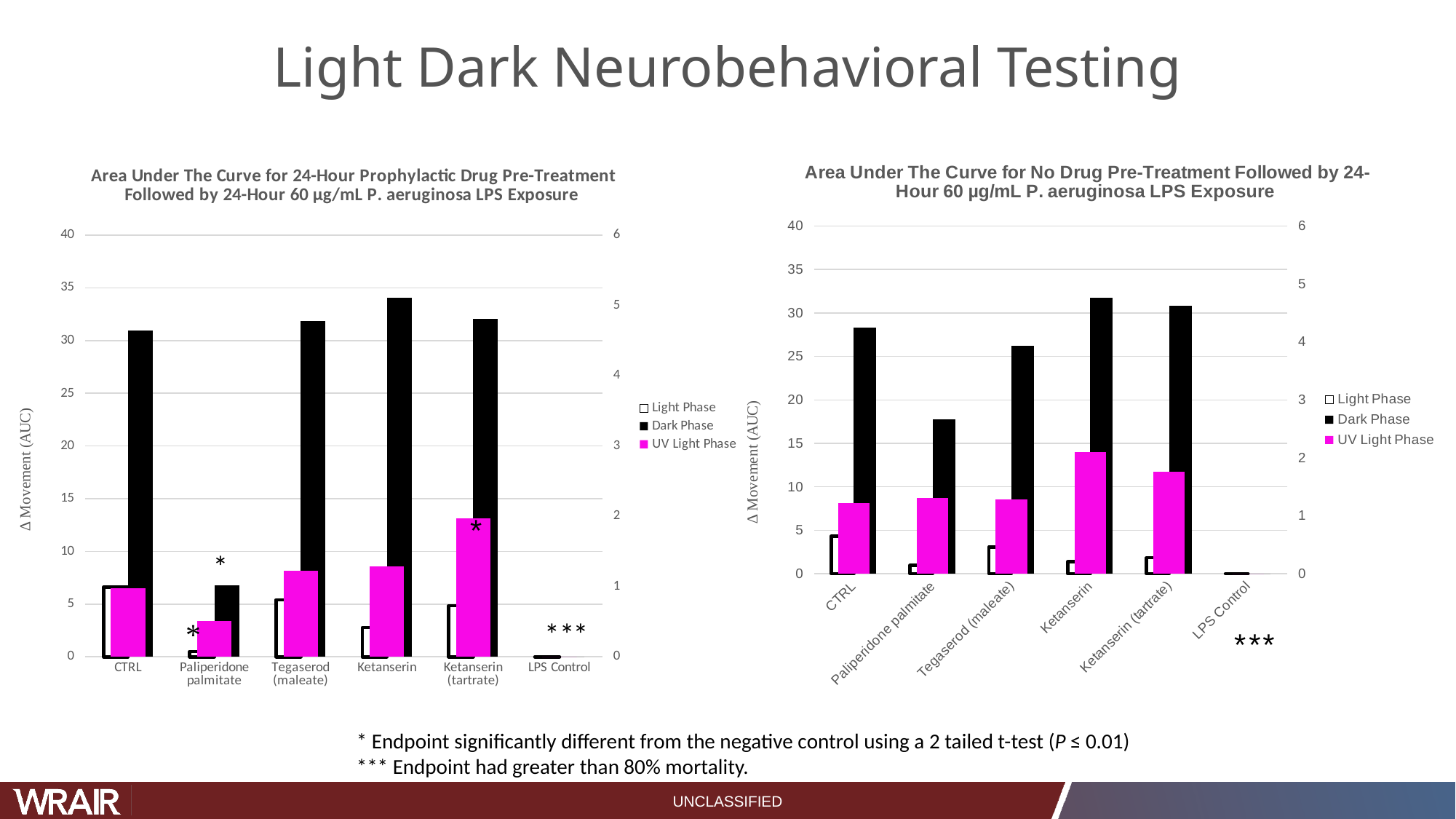

#### Light Dark Neurobehavioral Testing
###### Chart: Area Under The Curve for No Drug Pre-Treatment Followed by 24-Hour 60 µg/mL P. aeruginosa LPS Exposure
| Category | Light Phase | Dark Phase | UV Light Phase |
|---|---|---|---|
| CTRL | 4.315901327499997 | 28.337531807500003 | 1.21340459875 |
| Paliperidone palmitate | 0.9506089987499998 | 17.75298625250001 | 1.3118362834375001 |
| Tegaserod (maleate) | 3.089890334999999 | 26.2279875171875 | 1.2823099075 |
| Ketanserin | 1.3900727718750001 | 31.715319262187492 | 2.0941527875 |
| Ketanserin (tartrate) | 1.8355339415625 | 30.826674751562507 | 1.7607371159375003 |
| LPS Control | 0.0 | 0.0 | 0.0 |
###### Chart: Area Under The Curve for 24-Hour Prophylactic Drug Pre-Treatment Followed by 24-Hour 60 µg/mL P. aeruginosa LPS Exposure
| Category | Light Phase | Dark Phase | UV Light Phase |
|---|---|---|---|
| CTRL | 6.6135176559375015 | 30.948379440625015 | 0.9803878859375 |
| Paliperidone palmitate | 0.49399300968750004 | 6.779655204374996 | 0.5140845384375 |
| Tegaserod (maleate) | 5.401478088749998 | 31.879722512500006 | 1.2199472731249998 |
| Ketanserin | 2.7717947128124996 | 34.082229806250005 | 1.2856456115625001 |
| Ketanserin (tartrate) | 4.833485269374998 | 32.08348448531249 | 1.9724139021875 |
| LPS Control | 0.0 | 0.0 | 0.0 |*
* Endpoint significantly different from the negative control using a 2 tailed t-test (P ≤ 0.01)
*** Endpoint had greater than 80% mortality.
UNCLASSIFIED

### Slide 15
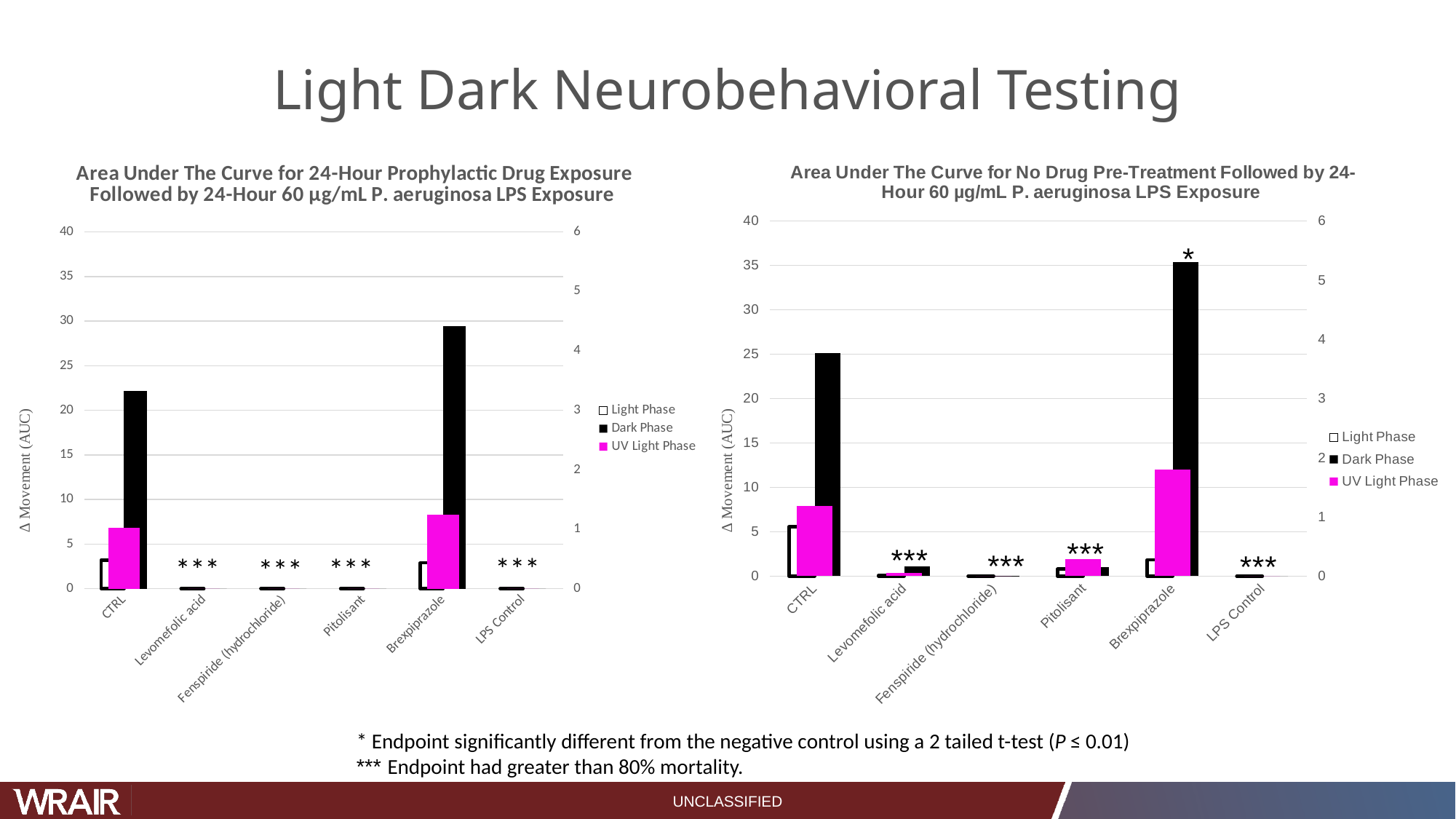

#### Light Dark Neurobehavioral Testing
###### Chart: Area Under The Curve for No Drug Pre-Treatment Followed by 24-Hour 60 µg/mL P. aeruginosa LPS Exposure
| Category | Light Phase | Dark Phase | UV Light Phase |
|---|---|---|---|
| CTRL | 5.569180865312498 | 25.096302549375004 | 1.186500865625 |
| Levomefolic acid | 0.09981209062499999 | 1.0697909868750002 | 0.060259059375000014 |
| Fenspiride (hydrochloride) | 0.0 | 0.0034823156250000003 | 0.0 |
| Pitolisant | 0.7885305262499996 | 1.0530600706249995 | 0.28755430843749996 |
| Brexpiprazole | 1.8320926575 | 35.3505170371875 | 1.79939065875 |
| LPS Control | 0.0 | 0.0 | 0.0 |
###### Chart: Area Under The Curve for 24-Hour Prophylactic Drug Exposure Followed by 24-Hour 60 µg/mL P. aeruginosa LPS Exposure
| Category | Light Phase | Dark Phase | UV Light Phase |
|---|---|---|---|
| CTRL | 3.21504220625 | 22.188215351562505 | 1.0205000421875001 |
| Levomefolic acid | 0.0 | 0.0 | 0.0 |
| Fenspiride (hydrochloride) | 0.0 | 0.0 | 0.0 |
| Pitolisant | 0.0 | 0.0 | 0.0 |
| Brexpiprazole | 2.8937779153124987 | 29.4673959021875 | 1.2384465053125 |
| LPS Control | 0.0 | 0.0 | 0.0 |* Endpoint significantly different from the negative control using a 2 tailed t-test (P ≤ 0.01)
*** Endpoint had greater than 80% mortality.
UNCLASSIFIED
